# Latent Transforming Growth Factor β Binding Protein 3 Controls Adipogenesis

**DOI:** 10.1101/2022.07.10.499446

**Authors:** Karan Singh, Nalani Sachan, Taylor Ene, Branka Dabovic, Daniel Rifkin

## Abstract

Transforming growth factor-beta (TGFβ) is released from cells as part of a trimeric latent complex consisting of TGFβ, the TGFβ propeptides, and either a latent TGFβ binding protein (LTBP) or glycoprotein-A repetitions predominant (GARP) protein. LTBP1 and 3 modulate latent TGFβ function with respect to secretion, matrix localization, and activation and, therefore, are vital for the proper function of the cytokine in a number of tissues. TGFβ modulates stem cell differentiation into adipocytes (adipogenesis), but the potential role of LTBPs in this process has not been studied. We observed that 72 h post adipogenesis initiation *Ltbp1*, *2*, and *4* expression levels decrease by 74-84%, whereas *Ltbp3* expression levels remain constant during adipogenesis. We found that LTBP3 silencing in C3H/10T1/2 cells reduced adipogenesis, as measured by the percentage of cells with lipid vesicles and the expression of the transcription factor peroxisome proliferator-activated receptor gamma (PPARγ). Lentiviral mediated expression of an *Ltbp3* mRNA resistant to siRNA targeting rescued the phenotype, validating siRNA specificity. Knockdown (KD) of *Ltbp3* expression in 3T3-L1, M2, and primary bone marrow stromal cells (BMSC) indicated a similar requirement for *Ltbp3*. Epididymal and inguinal white adipose tissue fat pad weights of *Ltbp3^-/-^* mice were reduced by 62% and 57%, respectively, compared to wild-type mice. Inhibition of adipogenic differentiation upon LTBP3 loss is mediated by TGFβ, as TGFβ neutralizing antibody and TGFβ receptor I kinase blockade rescue the LTBP3 KD phenotype. These results indicate that LTBP3 has a TGFβ-dependent function in adipogenesis both in vitro and in vivo.

**Significance:** Understanding the control of mesenchymal stem cell fate is crucial for the potential use of these cells for regenerative medicine.

**Highlights:**

- Latent TGFβ binding protein 3 (LTBP3) is required for adipogenesis
- LTBP3 mediates TGFβ levels in adipogenesis
- Loss of LTBP3 results in enhanced rather than decreased levels of active TGFβ

## Introduction

The cytokine TGFβ has multiple contextual effects on a variety of cell types during development and in adulthood. Loss of TGFβ results in abnormal bone formation, impaired lung development, inflammation, vascular defects, and cancer [1,2,3,4]. TGFβ activity is controlled both at the level of gene expression and receptor binding, as well as at the level of growth factor accessibility. Unlike most growth factors or cytokines, mature TGFβ is released from cells as part of an inactive complex in which the mature TGFβ homodimer remains non-covalently associated with its cleaved propeptide dimer [5]. Within this small latent complex (SLC), TGFβ cannot engage with its signaling receptor because the propeptides shield the receptor binding regions of the ligand. The propeptides are themselves disulfide-bonded either to one of three LTBPs, which are secretory proteins, or to GARP or to a leucine-rich-repeat-containing protein 32 (LRRC32), which are transmembrane proteins [6,7,8,9]. The trimeric complex of TGFβ, propeptide, and LTBP/GARP forms the large latent complex (LLC). LTBP, GARP and LRRC32 focus the LLC within extracellular space facilitating TGFβ activation by integrins, proteases, or shear. Formation of the LLC is necessary for proper TGFβ function, as prevention of LLC formation by mutation of the binding cysteine residues in either the propeptide or the LTBP blocks latent TGFβ activation [3, 10].

Three different LTBPs (LTBP1, 3, or 4) can bind covalently to the TGFβ propeptides [11,12,13,14]. LTBP1 and 3 avidly complex with all three isoforms of TGFβ, whereas LTBP4 binds only TGFβ1 and does so poorly. LTBP2 does not bind to any SLC isoform [10, 13]. The ability of multiple TGFβ isoforms to bind to two or three different LTBPs yields a combinatorial complexity associated with the ability of TGFβ to perform a plethora of functions, as individual LTBPs may have specific extracellular sites of deposition, unique expression patterns, or differential availability to activators of the latent complex.

One process in which LTBPs might have an important role is in the orchestration of stem cell differentiation. The local environment (niche) is important in directing stem cell differentiation, yet the identification of the extracellular components of the niche that participate in stem cell maintenance and developmental choice have not been well described. Within the mesenchymal stem cell niche, TGFβ acts as a suppressor of both adipo- and osteogenesis [15,16,17]. However, the activation of latent TGFβ and specifically the role of LTBPs in modulating TGFβ availability in this process within the niche have not been addressed. To interrogate the function of the LTBPs and, by inference TGFβ, in adipogenesis, we examined the LTBP requirements for cultured multipotent mesenchymal cell differentiation to adipocytes. We found that the elimination of LTBP3 in vitro and in vivo impedes adipocyte formation in a TGFβ-dependent manner.

## Results

### Effects of LTBP3 suppression

We examined LTBP levels during differentiation of mouse C3H/10T1/2 cells (10T1/2 cells) to gain insight into potential LTBP function in adipogenesis. 10T1/2 cells, when placed in culture medium that promotes adipogenesis, adopt an adipocyte phenotype that can be monitored by the expression of adipocyte-associated transcription factors, such as *Pparγ* and CCAAT/enhancer-binding protein-alpha (*Cebpα*) and by the accumulation of Oil Red O (ORO) positive lipid vesicles. To characterize the LTBP repertoire of 10T1/2 cells during adipogenesis, we initially exposed cells to adipocyte differentiation medium and quantified the transcript levels of the four *Ltbps*. Under non-differentiation conditions 10T1/2 cells express transcripts for all four *Ltbp* genes (**Fig. 1A-D**), but by 72 h post adipogenesis initiation, *Ltbp1*, *2*, and *4* transcript levels are decreased by 74-84% and by 120 h by 89-97% (**Fig. 1A, B and D**), whereas *Ltbp3* transcript levels remained virtually unchanged (**Fig. 1C**). Transcript levels of *Pparγ* significantly increased over the course of the experiment (**Fig. 1E**). The sustained expression of *Ltbp3* suggested a potential continuing role for this TGFβ carrier in the differentiation process.

We probed the role of LTBP3 during 10T1/2 cell differentiation by treating cells with a siRNA (siLtbp3-4) designed to facilitate *Ltbp3* mRNA degradation. siLtbp3-4 pre-treatment for 48 h diminished LTBP3 protein levels by over 64% for at least 7 days (**SI Fig. 1C and D**). The long-term (5 days) suppression of LTBP3 protein levels permitted us to treat cells with siLtbp3-4, wait 48 h to eliminate existing *Ltbp3* mRNA, initiate differentiation, and measure levels of adipocyte markers over the subsequent 3-5-day period. In addition to the decrease in *Pparγ* observed upon *Ltbp3* KD, we also detected a concordant decrease in the expression of the early transcription factors sterol regulatory element-binding factor-1 (*Srebf1*), CCAAT/enhancer binding protein-delta *(Cebpδ)*, the adipogenic master transcription factors *Pparγ* and *Cebpα,* as well as fatty acid synthase (*Fasn*); all of which are associated with adipogenesis (**Fig. 1G, I**-**L**). There was only a slight decrease in the expression level of the early transcription factor *Cebpβ* (**Fig.1H**). In addition, there was a significant decrease (80%) in cells containing lipid vesicles (**SI Fig. 1A and B**), indicating the acquisition of an adipocyte phenotype. These results indicate the participation of LTBP3 in the initiation of the adipogenic state.

**Fig. 1.**
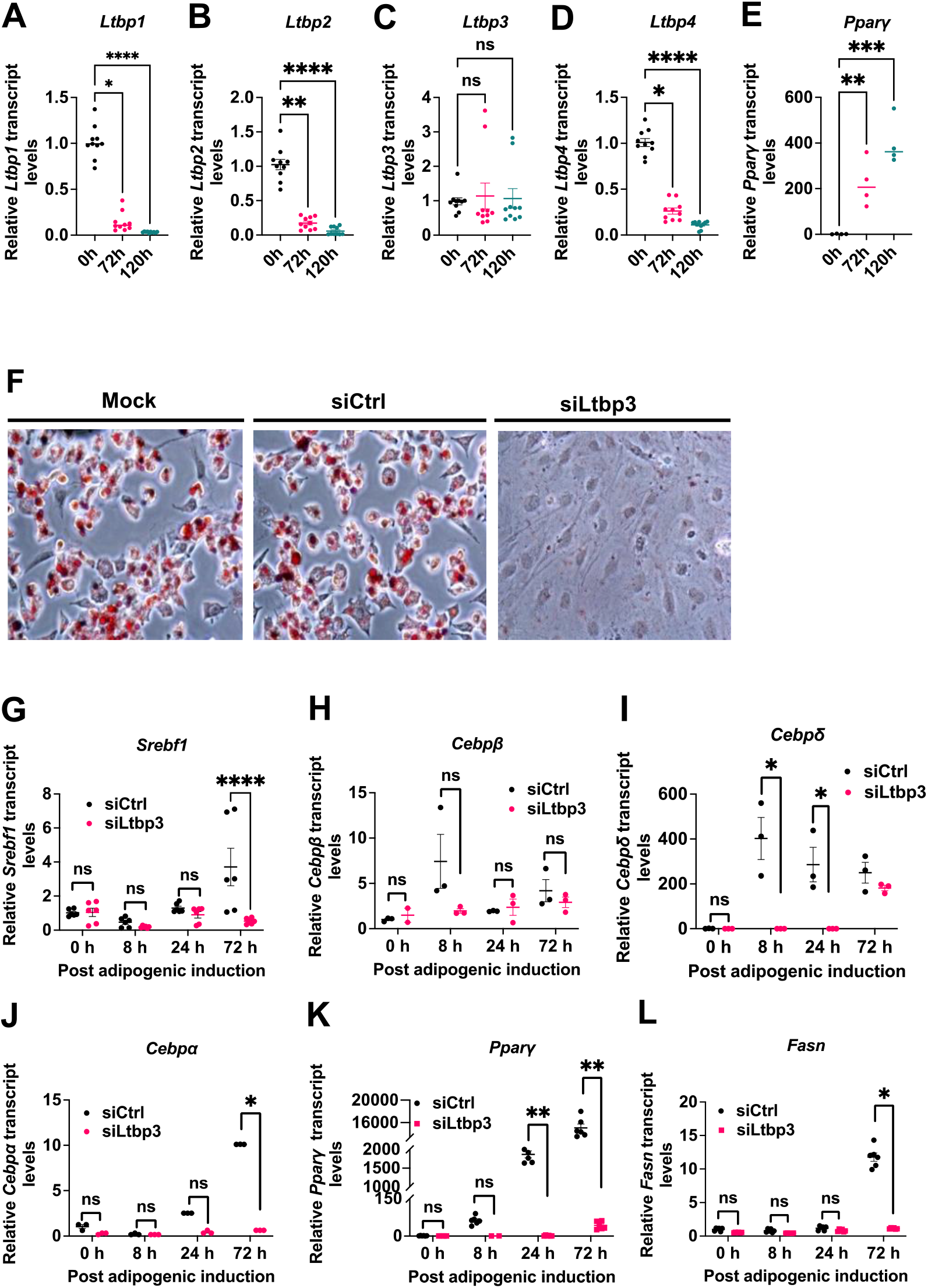
LTBP3 regulates adipogenesis in vitro. **(A-E)** Relative transcript levels of *Ltbp1*, *Ltbp2*, *Ltbp3*, *Ltbp4*, and *Pparγ* in 10T1/2 cells. Levels of the four different *Ltbps* and *Pparγ* transcripts in 10T1/2 cells treated with adipogenic media for 0, 24, 72, and 120 h were measured by qRT-PCR as described in Methods. qRT-PCR values were normalized to beta-2 microglobulin (*B2m*) and plotted relative to siCtrl. Data represent the average of four independent experiments. (**F**) Representative images illustrating accumulation of ORO-stained lipid vesicles or droplets in 10T1/2 cells treated with mock, siCtrl, or siLtbp3-4 RNAs and exposed to adipogenic media for 5 days. Scale bar, 100 µm. **(G**-**L**) Relative transcript levels of *Srebf1*, *Cebpβ, Cebpδ*, *Pparγ, Cebpα,* and *Fasn* measured at 0, 8, 24, and 72 h after adipogenic induction with 10T1/2 cells treated with siCtrl or siLtbp3-4 for 48 h. qRT-PCR values were normalized to *B2m* and plotted relative to siCtrl. Each value is the average of 3 independent experiments. For figures **A**-**D**, there were 10 technical replicas normalized and pooled from four experiments and each replica was analyzed twice by qRT-PCR. For figure **E,** the data represent a total of 4 samples for each condition normalized and pooled from four experiments (1 for each). Figure **F** is representative of one of 3 independent experiments in which there was 1 technical replica. For figures **G**, **K**, and **L** each data point represents two technical replicas from three experiments. Each technical replica was analyzed two times by qRT-PCR. For figures **H**, **I**, and **J**, each data point represents one technical replica from three experiments. Each technical replica was analyzed once with qRT-PCR. Statistical significance was evaluated by Nonparametric, Kruskal-Wallis test with Dunn’s multiple comparisons test (**Fig**. **A**-**E**), or two-way mixed model analysis of variance (ANOVA) using time and treatment as fixed factors with Tukey’s multiple comparisons test (**Fig. G**-**L**). Data are represented as mean ± SEM. *p < 0.05, **p < 0.01, ***p < 0.001, ****p < 0.0001.

### Specificity of *Ltbp3* knockdown

We validated that the effect of siLtbp3-4 is specific for *Ltbp3* by several approaches. First, four different siRNAs (siLtbp3-1-4), three (siLTBP3-1, 2, and 3) targeted to unique sequences in the *Ltbp* 3” UTR and one (siLTBP3-4) targeted to the coding sequence in *Ltbp3* exons 13-14, decreased both *Ltbp3* expression and cell differentiation, as monitored by *Pparγ* expression (**Fig. 2A and B**). siLtbp3-3 was not quite as effective as siLtbp3-1, 2, or 4 and there appeared to be a relationship between the degree of *Ltbp3* suppression and *Pparγ* expression (**Fig. 2A and B**).

Second, we prepared a rescue lentivirus vector containing a cloned *Ltbp3* missing the 3’ UTR. This permitted us to employ siLtbp3-2, which targets the 3’ UTR, to eliminate specifically the endogenous *Ltbp3* transcripts, but which would not recognize the lentiviral vector encoded *Ltbp3* transcripts, as they are derived from a cDNA. We validated the occurrence of expressed LTBP3 in 10T1/2 cells, as detected by immunofluorescence, as well as by measuring LTBP3 protein as well as transcripts (**SI Fig. 2A-C**). Cells transduced with the rescue LTBP3 lentivirus, treated with siLtbp3-2, and placed in differentiation medium continued to express *Ltbp3* mRNA (**Fig. 2C**) and protein (**SI Fig. 2D, E**). These cells were resistant to the interference of adipogenesis, as measured by *Pparγ* and *Cebpα* expression (**Fig. 2D and E**), the percentage of cells with lipid vesicles (**Fig. 2F and G**), and PPARγ protein levels (**SI 2Fig. D and E**). The expression level of the rescue *Ltbp3* mRNA was consistently ∼2 fold higher than that of the endogenous transcripts of *Ltbp3* gene (**Fig. 2C**). The reason for this is unclear but could relate to the lack of the 3’ UTR or to the viral titer. This question was not pursued. As expected, cells transduced with a lentivirus expressing *Ltbp3* transcripts were not rescued with respect to *Pparγ* expression when treated with siLtbp3-4, which recognizes a coding region of *Ltbp3* mRNA (**SI Fig. 2F and G**). These results indicate that the effect of siLtbp3-2 mediated *Ltbp3* KD is not the result of an off-target activity.

**Fig. 2.**
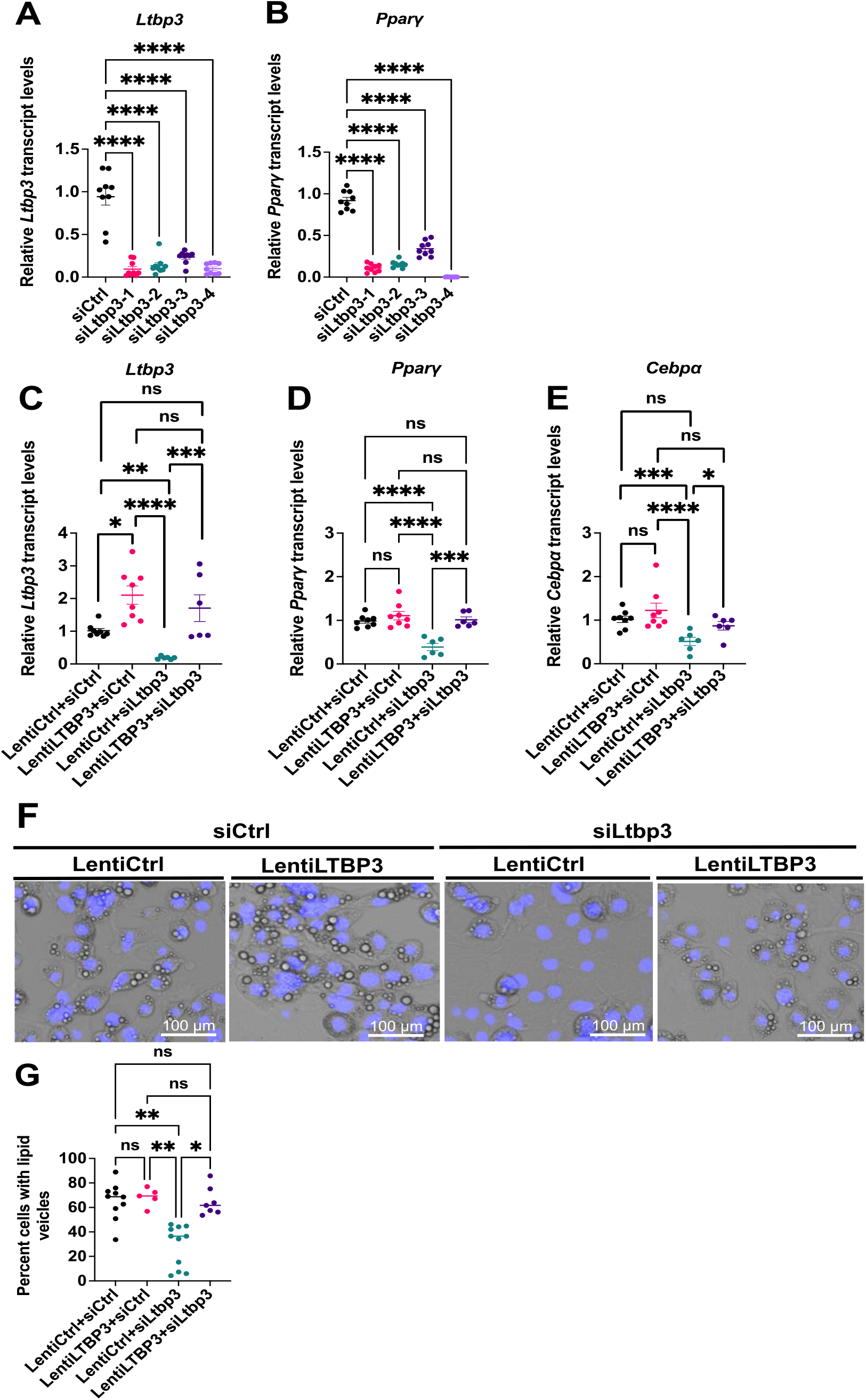
*Ltbp3* knockdown specificity. **A)** and **B)** Effect of different *Ltbp3* siRNAs on adipogenesis. Relative mRNA levels of *Ltbp3* and *Pparγ* were measured in 10T1/2 cells treated with siCtrl or siLtbp3-l, 2, 3, or 4, maintained for two days in basal media, and incubated for three days in adipogenic medium. qRT-PCR values were normalized to *B2m* and plotted relative to the siCtrl. Data represent the means of 3 independent experiments with 3 technical replicas for each sample, and each technical replica was analyzed 2 times by qRT-PCR. (**C-E)** Relative mRNA levels of *Ltbp3* (**C**)*, Pparγ* (**D**), *and Cebpα* (**E**) in LentiCtrl and LentiLTBP3 cells treated with siCtrl or siLtbp3-2 for 48 h followed by 72 h of adipogenic induction. Data represent the means of 3 independent experiments with 1-3 technical replicas for each group, and each replica was analyzed 2 times by qRT-PCR. (**F)** Lipid vesicles in cells rescued by lentivirus mediated expression of *Ltbp3*. Images show birefringent lipid vesicles and blue DAPI stained nuclei in LentiCtrl and LentiLTBP3 cells treated with siCtrl or siLtbp3-2 for 48 h followed by 120 h incubation in adipogenic medium. Scale bar, 100 µm. (**G)** Quantification of cells with lipid droplets from panel **F.** For **Fig. F** representative images are from one of three independent experiments with two-four technical replicas and for **Fig. G** data are the mean of two independent experiments with 2,000-4,000 cells counted in each replica. Statistical significance was evaluated by one-way ANOVA with Dunnett’s multiple comparison (**Fig. A** and **B**) or nonparametric, Kruskal-Wallis test with Dunn’s multiple comparisons test. (**Fig. C**-**E** and **G**). Data are represented as mean ± SEM. *p < 0.05, **p < 0.01, ***p < 0.001, ****p < 0.0001.

Third, we tested whether the restraint of adipogenesis by siLtbp3-2 was restricted to 10T1/2 cells. We monitored the result of *Ltbp3* loss on differentiation of the mouse preadipocyte line 3T3-L1, the mouse bone marrow-derived mesenchymal cell line M2, and the mouse bone marrow-derived stromal primary cell (BMSC). KD of *Ltbp3* transcripts in all three cell types resulted in impaired *Pparγ* expression (**Fig. 3A-H**). These data plus results with *Ltbp3* KD of primary mouse BMSC indicate that the *Ltbp3* requirement for adipogenesis is not unique to 10T1/2 cells.

**Fig. 3.**
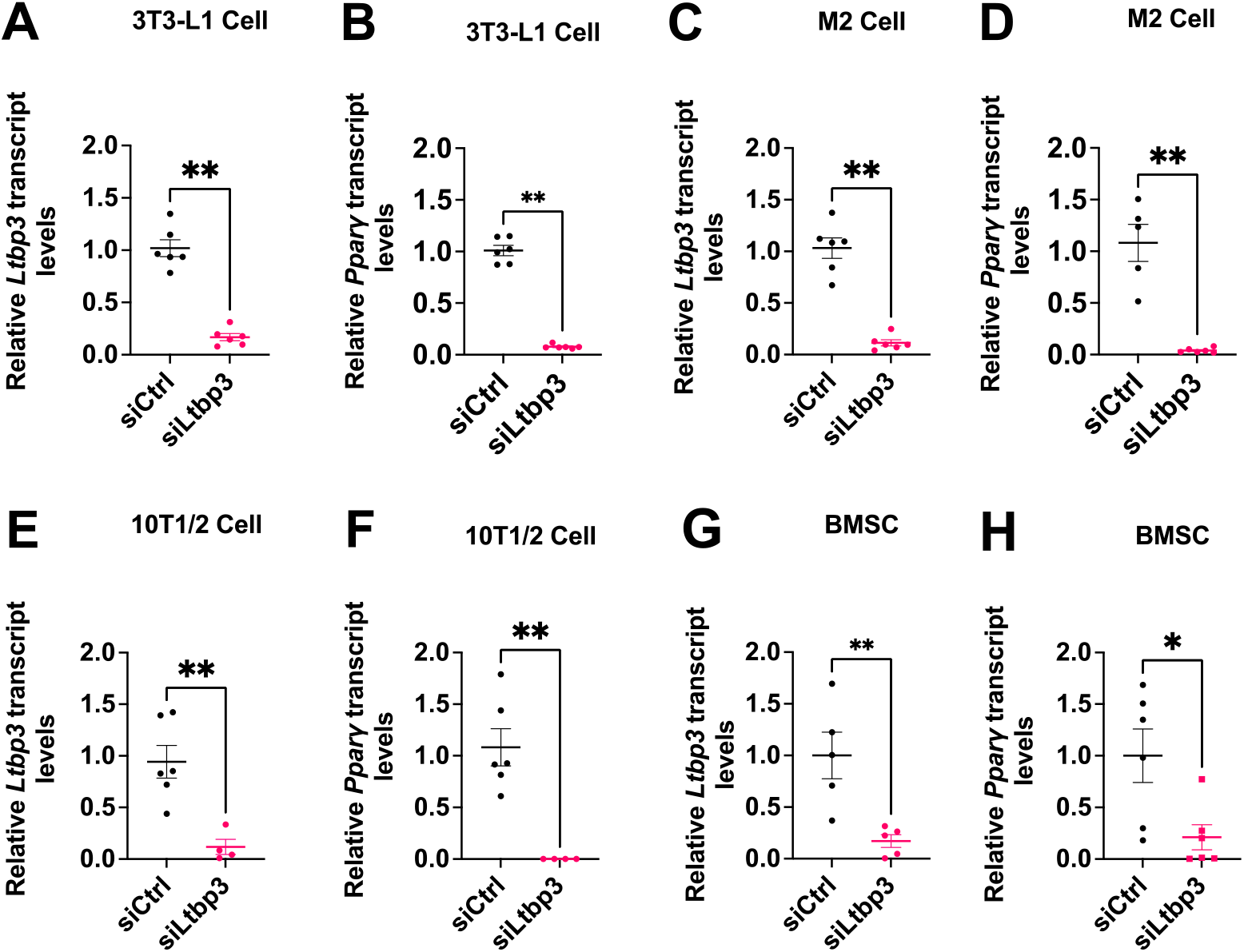
Effect of *Ltbp3* downregulation in 3T3-L1, 10T1/2, M2, and BMSC. (**A**-**H)** Relative transcript levels of *Ltbp3* and *Pparγ* measured at 96 h post siCtrl or siLtbp3-2 treatment (48 h in basal media followed by 48 h of adipogenic stimulation except for BMSC, BMSC were exposed to adipogenic media for 72 h) with 3T3-L1 (**A** and **B**), M2 (**C** and **D**), 10T1/2 (**E** and **F**), and BMSC (**G** and **H**) cells. For **Fig. A-F,** data are the mean ± SEM of 3 independent experiments with one-three technical replicas for each experiment and treatment condition. Each technical replica was analyzed two times by qRT-PCR. For experiments **G** and **H**, there were 5-6 animals per group and the samples were each assayed in duplicate by qRT-PCR. Statistical significance was evaluated by nonparametric, Mann-Whitney U test (**A**-**H)**. *p < 0.05, **p < 0.01, ***p < 0.001, ****p < 0.0001.

### Effect of in vivo loss of *Ltbp3*

To probe if *Ltbp3* loss in vivo yields decreased fat accumulation, we performed DEXA scans on wild-type (*WT)* and *Ltbp3* null mice to measure body composition. DEXA scans revealed that male *Ltbp3* null mice had a ∼30% decrease in fat mass and a corresponding ∼13% increase of lean mass by 18 weeks of age (**Fig. 4A and B**). We next visualized the white fat depots to see if specific body fat deposits differed between *WT* and *Ltbp3* null mice. A clear contrast was apparent in the size of inguinal (iWAT) and epididymal (eWAT) subcutaneous fat pads between males of the two genotypes (**Fig. 4C**). As reported previously [18], there was also a significant loss of body weight in the *Ltbp3^-/-^* animals compared to *Wt* counterparts (**Fig. 4D**). When we dissected animals, weighed and normalized fat pads to body weight, we found a 62% loss of eWAT and a 57% loss of iWAT in *Ltbp3* null compared to *WT* animals (**Fig. 4E and F**). However, we observed no significant difference in liver, a potential site for fat deposition, weights between *WT* and *Ltbp3^-/-^* animals (**Fig. 4G**). The fat mass loss was not as dramatic in female mice, which had fat mass losses of less than 18% and a corresponding ∼8% increase of lean mass by age of 18 weeks (**Fig. 4H and I**). Female mice of the two genotypes also displayed no significant differences in body weight, eWAT, iWAT, or liver weight (**Fig. 4J-M**). The decreased amount of white fat in *Ltbp3^-/-^* animals was consistent with the in vitro findings described above. However, *Ltbp3* loss might have indirect or systemic effects that modulate fat accumulation in the animal. Therefore, we evaluated the ability of primary *WT* BMSC to differentiate into adipocytes. When we treated freshly isolated *WT* BMSC with siLtbp3-2 and induced them to differentiate, we observed a significant decrease in *Ltbp3* transcripts (**Fig. 3G**) and protein (**SI Fig. 3A and B**), the degree of differentiation measured by *Pparγ* mRNA (**Fig. 3H**) and protein levels (**SI Fig. 3A and C**), and the number of cells with lipid vesicles (**SI Fig. 3D and E**). We observed a similar inhibition of adipogenesis as measured by the number of cells containing lipid vesicles (**SI Fig. 3F and G**) and *Pparγ* mRNA transcripts (**Fig. 4N**) using BMSC derived from *Ltbp3*^-/-^ bone marrow (**Fig. 4O**). Thus, *Ltbp3* loss mediated by either siRNA mediated KD or gene targeting impairs adipogenesis.

**Fig. 4.**
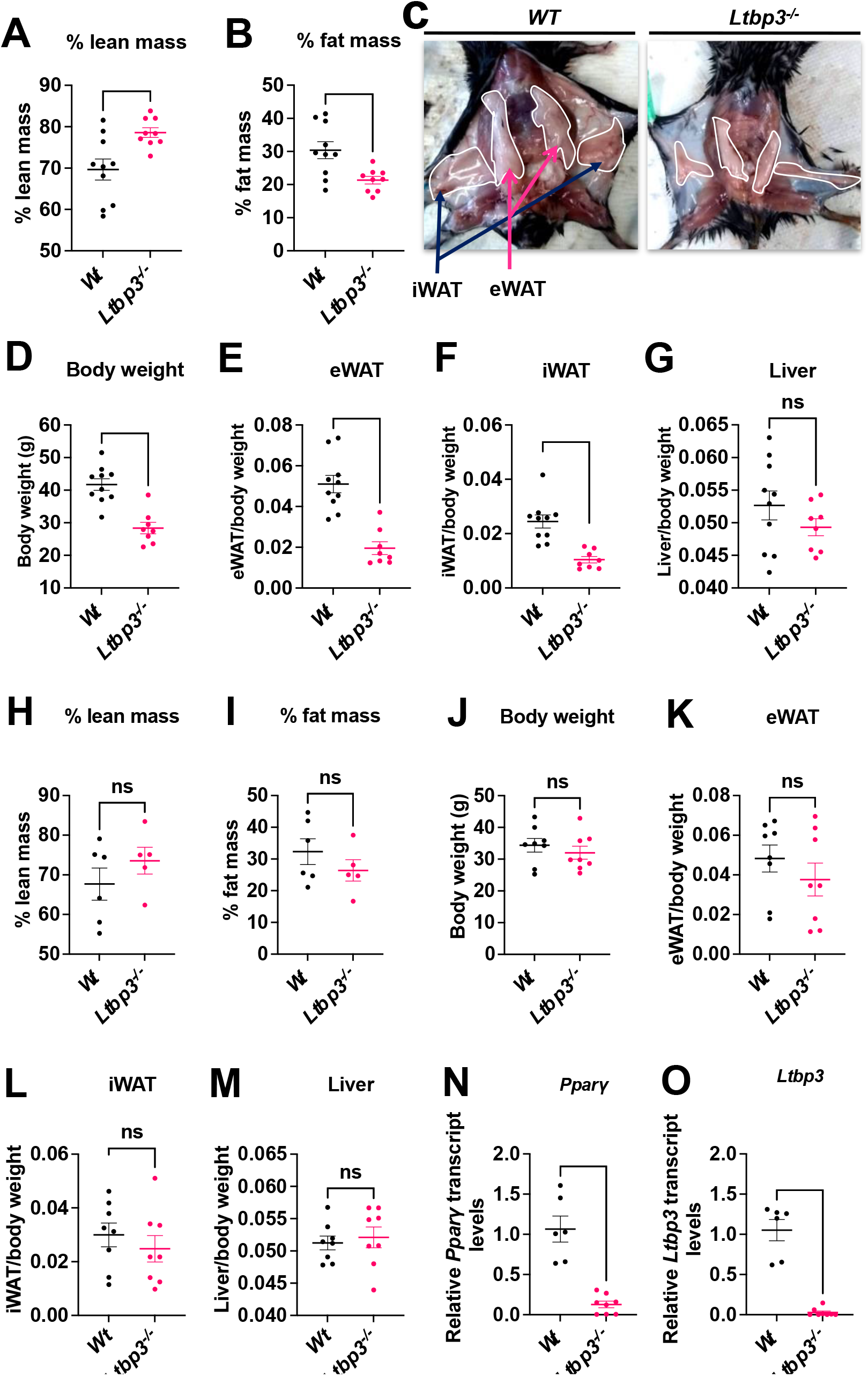
Reduced adipose tissue in *Ltbp3^-/-^* mice. (**A** and **B**) DEXA scans for body composition of 18-week-old male mice. (**A**) percent lean mass, (**B**) percent fat mass. (**C)** Representative images of the fat depots of 18-week-old *WT* and *Ltbp3^-/-^*male mice; epididymal white adipose tissue (eWAT) and inguinal white adipose tissue (iWAT). (**D)** Body weights of 18-week-old *WT* and *Ltbp3^-/-^* male mice. **(E-G)** Relative weights of eWAT, iWAT, and liver normalized to body weights of 18-week-old WT or *Ltbp3^-/-^* male mice. (**H** and **I**) DEXA scans for body composition of 18-week-old female mice. **H** percent lean mass, **I** percent fat mass. (**J)** Body weights of 18-week-old *WT* and *Ltbp3^-/-^* female mice. **(K-M)** Relative weights of eWAT, iWAT, and liver normalized to body weights of 18-week-old WT or *Ltbp3^-/-^* female mice. **(N** and **O)** Relative transcript levels of *Pparγ* and *Ltbp3* measured at 120 h post differentiation treatment (48 h in basal media followed by 72 h of adipogenic stimulation) of *Wt* and *Ltbp3*^-/-^ BMSC. For experiments in parts **Fig. 4A-B, D-G and J-M** there were 8-10 animals per group, and for part **Fig. 4H-I** there were 5-6 animals per group. For experiments **N** and **O**, there were 6-8 animals per group and the samples were each assayed in duplicate for the qRT-PCR. Data are represented as mean ± SEM. Statistical significance was confirmed by using nonparametric, Mann-Whitney U test (**Fig. A**-**B** and **D**-**O**). p < 0.05, **p < 0.01, ***p < 0.001, ****p < 0.0001.

### Requirement for TGFβ

LTBP null phenotypes in both humans and mice have been related to decreased TGFβ signaling, consistent with the hypothesis that LTBPs are critical mediators of TGFβ function. However, decreased TGFβ resulting from the loss of LTBP3 is unlikely to account for impaired adipogenesis, since TGFβ is a known suppressor of adipocyte differentiation. Thus, LTBP3 loss and consequent decrease in active TGFβ should enhance differentiation. Nevertheless, we tested the potential involvement of TGFβ in our system by several approaches. We initially monitored the effect of TGFβ supplementation on *Ltbp3* KD 10T1/2 cells (**SI Fig. 4A and B**). Addition of TGFβ1 yielded cultures with enhanced impairment of differentiation compared to those with *Ltbp3* KD alone, indicating, as predicted, that TGFβ loss was not responsible for the inhibition of adipogenesis (**SI Fig. 4A and B**).

We next examined the level of TGFβ signaling in control and LTBP3 KD cells cultures by monitoring phospho-SMAD3 (p-SMAD3) levels, as p-SMAD3 can be used as a surrogate marker for TGFβ activity. Indeed, we observed an increase (∼2.3-fold) in p-SMAD3 after treatment with the siLtbp3-2 (**Fig. 5A and B**), implying that there was an increase in active TGFβ after inhibition of LTBP3 production. We reasoned that if excess TGFβ produced after *Ltbp3* KD was responsible for the inhibition of adipogenesis, prevention of TGFβ signaling either with an inhibitor to the TGFβ receptor kinase or a neutralizing antibody should overcome the KD effect. Addition of the low molecular weight TGFβ type I receptor kinase (ALK5) inhibitor (SB431542) effectively decreased p-SMAD3 levels (**Fig. 5A**), and rescued the *Ltbp3* KD phenotype as monitored by recovery of PPARγ protein (**Fig. 5C and D**) and transcript levels (**SI Fig. 4C and D**), as well as the number of cells with lipid vesicles (**Fig. 5E and F**). As expected, there was no effect of SB431542 treatment on LTBP3 accumulation (**Fig. 5A-D, SI Fig, 4C**).

**Fig. 5.**
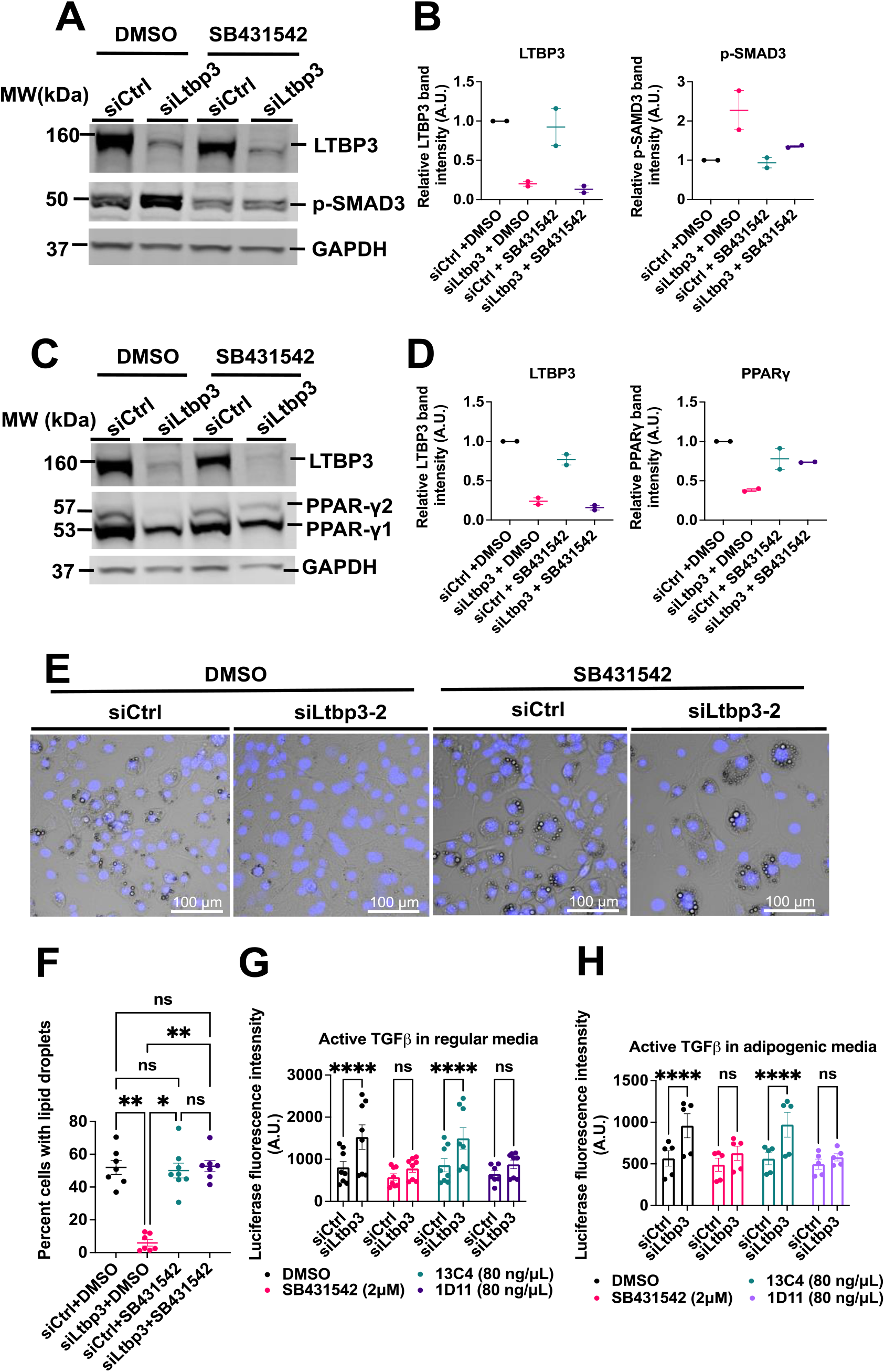
Heightened TGFβ inhibits adipogenesis. (**A**-**F**) TGFβ receptor kinase inhibitor reverses the LTBP3 KD effect. 10T1/2 cells were treated with SB431542 as described in Methods. (**A-D**) Cell lysates were analyzed by immunoblotting with antibodies to LTBP3, PPARγ, p-SMAD3, and GAPDH. The immunoblot is representative of one of two independent experiments. There was 1 technical replica for each treatment condition. (**E**) Images showing accumulation of lipid droplets in 10T1/2 cells treated with siCtrl or siLtbp3-2 and exposed to TGFβ receptor1 kinase (ALK5) inhibitor as described in Methods. Representative images are shown from one of three independent experiments. For each experiment there was 1 technical replica and 4-5 fields were counted for each replica (1,800-4,000 cells for each replica). Scale bars, 100 μm. **(F)** Quantification of percent cells with lipid droplets for panel **E**. (**G** and **H**) Active TGFβ produced by siCtrl or siLtbp3-2 treated 10T1/2 cells was determined using a luciferase reporter cell assay. 10T1/2 cells were treated as described in Methods. The **Fig. G** represents the values from three independent experiments and **Fig. H** from two independent experiments. Each treatment had 2-3 technical replicas. Statistical significance of **Fig. F** was evaluated by the using nonparametric, Kruskal-Wallis test with Dunn’s multiple comparisons test. Data are represented as means ± SEM. p < 0.05, **p < 0.01, ***p < 0.001, ****p < 0.0001.

We also attempted to rescue the *Ltbp3* KD phenotype, by addition of an antibody that neutralizes all three isoforms of TGFβ to siCtrl or siLtbp3-2 treated cells. Although an isotype control antibody had no effect on LTBP3 or p-SMAD levels in siCtrl or siLtbp3-2 treated cells, the inclusion of an antibody that neutralized TGFβ resulted in a significant decrease in p-SMAD3 induction by siLtbp3-2 (**SI Fig. 5A and B**). In addition, when we treated cells with the TGFβ neutralizing antibody, there was an increase in PPARγ protein (**SI Fig. 5C and D).** Although the increase in PPARγ protein was modest, when we compared the numbers of siLtbp3-2 treated cells plus neutralizing antibody vs. cells treated with control antibody containing lipid vesicles, we observed that the recovery of the adipogenic state was highly significant (**SI Fig. 5E and F)**. The fact that the effect of antibody treatment was less than that observed with the kinase inhibitor may relate to the greater ease of the low molecular weight inhibitor, compared to the antibody, to reach its target.

These results are consistent with increased rather than decreased TGFβ. Therefore, we quantified the level of active TGFβ in cultures of siLtbp3-2 treated 10T1/2 cells in regular and adipogenic media. When cells were cultured in regular medium, the presence of the siLtbp3 clearly enhanced the level of TGFβ and the induction was blocked by both the TGFβ receptor 1 kinase inhibitor as well as the TGFβ specific neutralizing antibody (**Fig. 5G**). We observed a lower, but significant, increase in TGFβ under the different treatments when cells were cultured in the adipogenic medium (**Fig. 5H**). The decreased TGFβ levels in cultures with adipogenic medium was due to the effect of the dexamethasone in the adipogenic culture medium on the plasmid promoter used in the assay. Together these data indicate that increased TGFβ signaling is responsible for the inhibition of differentiation upon loss of LTBP3.

**Supplementary Figure 5.**
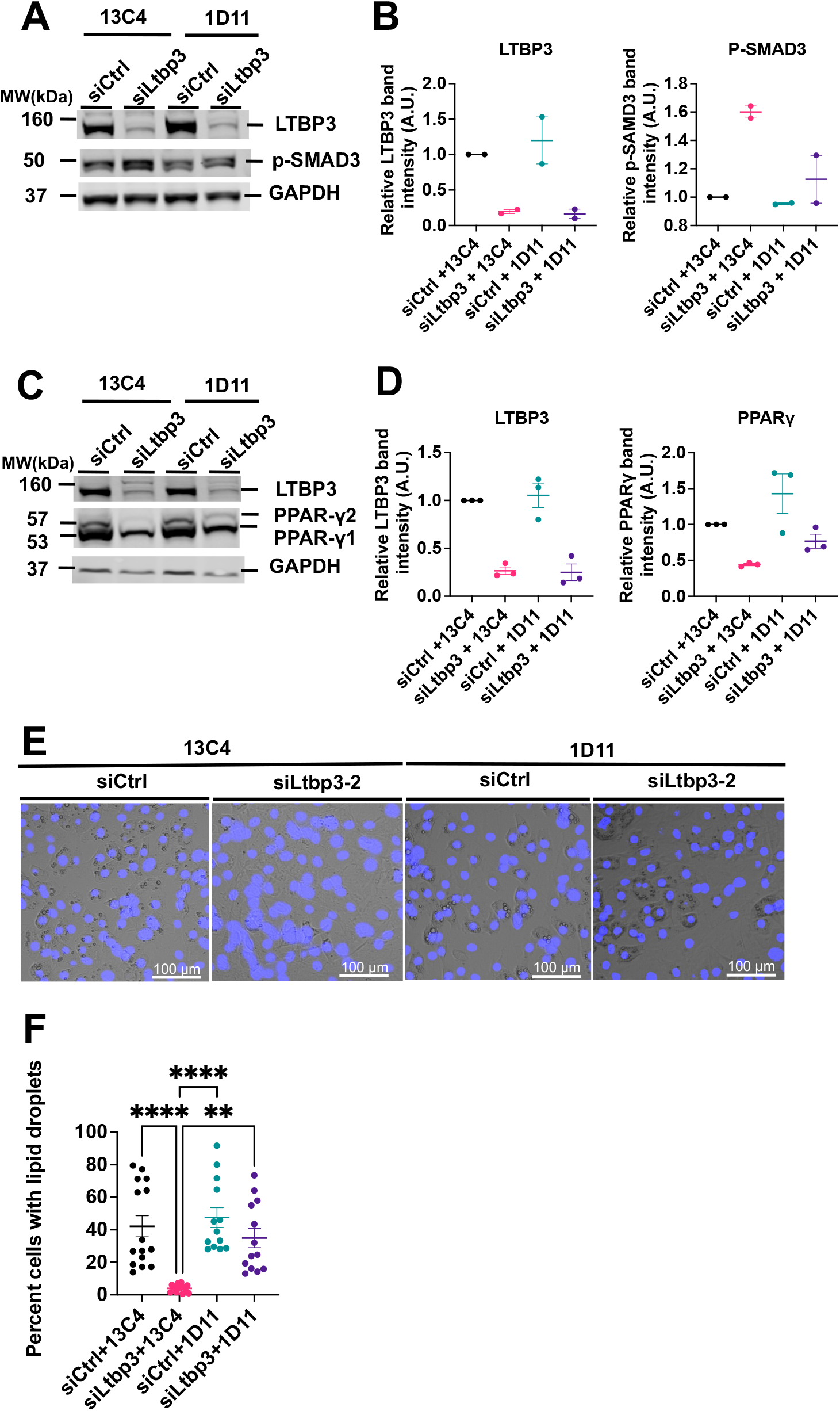
Antibody inhibition of TGFβ rescues adipogenesis in *Ltbp3* KD 10T1/2 cells. **(A)** Immunoblot illustrating the levels of LTBP3, p-SMAD3 and GAPDH in 10T1/2 cells treated with siCtrl or siLtbp3-2 for two days followed by 3 days of adipogenic differentiation media treatment with control (13C4) or specific TGFβ (1D11) neutralizing antibodies. The experiment was repeated twice. (**B**) Quantification of LTBP3 and p-SMAD3 from figure **A**. (**C**) Immunoblot illustrating the levels of LTBP3, PPARψ and GAPDH in 10T1/2 cells treated with siCtrl or siLtbp3-2 for two days followed by 3 days of adipogenic differentiation media treatment with control (13C4) or TGFβ specific (1D11) antibodies. The experiment was repeated three times. (**D**) Quantification of LTBP3 and PPARψ from figure **C**. (**E**) Representative images showing accumulation of lipid droplets (white vesicles) and DAPI stained nuclei (blue) in 10T1/2 cells treated with siCtrl or siLtbp3-2 for two days and exposed to control (13C4) or TGFβ specific (1D11) antibodies in adipogenic media for 5 days. The experiment was repeated three times. (**F**) Quantification of lipid loaded cells after differentiation was induced in the presence of control (13C4) or TGFβ specific (1D11) neutralizing antibodies. The number of cells with lipid droplets, as well as total number of cells, were counted manually. 4-5 random fields from each treatment group were counted for each experiment. Data were taken from three independent experiments. Scale bar, 100 µm. Statistical significance of **Fig. F** was evaluated by the using nonparametric, Kruskal-Wallis test with Dunn’s multiple comparisons test. Data are represented as means ± SEM. p < 0.05, **p < 0.01, ***p < 0.001, ****p < 0.0001.

## Discussion

The LTBPs are important for the effective secretion and localization of latent TGFβ into the extracellular matrix and are perceived to be crucial for the activation of the latent cytokine [19]. Here we present evidence that LTBP3 loss inhibits preadipocyte and mesenchymal stem cell adipogenesis, as measured by the impaired accumulation of lipid vesicles and by the decrease of specific transcription factor expression in both cultured and primary cells. We documented the specificity of the effect by rescuing siRNA mediated inhibition using a lentiviral vector expressing an *Ltbp3* mRNA resistant to siRNA mediated degradation, by blocking *Ltbp3* expression in four different cell types with consequently impaired adipogenesis, and by demonstrating that both *Ltbp3*^-/-^ cells and animals have impaired ability to form adipocytes. We rescued adipogenesis in LTBP3 KD cells by blocking TGFβ signaling either with a TGFβ neutralizing antibody or by inhibiting the TGFβ receptor kinase I indicating that active TGFβ is the effector molecule in LTBP3 silencing. Our studies are the first to identify an LTBP required for stem cell differentiation.

Our results are consistent with the known inhibitory effect of TGFβ on adipogenesis with both pre-adipocytes and bone derived mesenchymal cells [16,17, 20,21,22,23]. Additionally, Clouthier et al. documented that transgenic overexpression of human TGFβ1 in white adipose tissue hampered adipogenesis [24]. These earlier studies revealed that TGFβ signaling represses C/EBPβ and C/EBPδ functions by binding to activated Smad3 [16]. Similarly, Smaldone et al. [25] reported that heightened TGFβ signaling in cultured marrow cells from mouse limbs deficient in fibrillin-1 impaired adipogenesis, as measured by PPARγ expression. An interesting question is whether LTBP3 controls the differentiation of MSCs along lineages other than the adipogenic lineage or is there lineage specificity. Since TGFβ is known to inhibit chondrogenesis [26, 27] and osteogenesis [28], we expect that loss of LTBP3 will impair differentiation along these additional pathways.

Unexpectedly, we found a sexual dimorphism with regard to adiposity and LTBP3 loss with females exhibiting no statistically significant differences in the absence of LTBP3. Sexual dimorphism in fat accumulation has been described for a subset of mice with fibrillin1 mutations [29]. However, there appear to be no other publications indicating that loss of additional matrix components yields sex-specific differences in weight gain. This could, however, reflect the common practice of primarily focusing on the outcomes with male mice.

An interesting question is whether other LTBPs, for example LTBP1, might compensate or cooperate with LTBP3 during adipogenesis. Or conversely, is suppression of LTBP1, 2, and 4 required for proper adipogenesis? We have shown that the levels of expression of other LTBPs all decrease dramatically during the time period in which we have measured adipogenesis. However, we cannot rule out the re-expression at later times. It does appear that Ltbp3 null animals display decreased fat accumulation indicating that any compensation by a second LTBP is minimal. Moreover, genetic studies in mice have failed to reveal any compensation of the loss of LTBP3 in either lung or aorta when LTBP is missing in the presence of other LTBPs [30, 31].

The participation of LTBPs in stem cell differentiation has not been extensively studied. Koli et al. [32], who examined human mesenchymal stem cell osteogenic differentiation, and Gualandris et al. [33], who monitored mouse embryoid body differentiation, both concluded that differentiation required TGFβ and effective activation of latent TGFβ required the participation of an LTBP. Thus, the loss of an LTBP yielded diminished TGFβ levels. However, our findings indicate LTBP3 down regulation results in an increase, rather than decrease, in active TGFβ during adipogenesis; a finding more in concert with those of Smaldone et al. (vide supra) [25].

Several lines of experimentation have led to the concept that LTBPs support latent TGFβ activation and that the loss of LTBP1 or LTBP3 results in decreased active cytokine levels. Early investigations with antibodies to LTBP1 indicated that interference with LTBP1 reduced TGFβ activity [34, 35]. Subsequent phenotypic analysis of mice with LTBP1 and LTBP3 null mutations revealed pathologies also congruent with loss of TGFβ function [18,36,37]. The discovery that integrin binding to the TGFβ propeptide facilitated LLC activation suggested that force was necessary for latent TGFβ activation and that the immobilization of the SLC by covalent binding to an LTBP or GARP/LRRC32 provided the required traction [38,39,40]. The SLC crystal structure revealed how integrin pulling at one end of the latent complex would distort the propeptides and liberate active TGFβ [41, 42]. Finally, mice with a mutation in the TGFβ1 propeptide residue that binds to the LTBP or GARP/LRRC32 display phenotypes overlapping with those of TGFβ 1 null mice, implying a requirement for LLC formation to facilitate latent TGFβ1 activation [3]. Therefore, binding of the SLC to an LTBP is thought to be required to develop the tension necessary for integrin mediated activation of the latent complex. The phenotypes apparent after LTBP3 loss in mice, such as premature ossification of the synchondroses [18], amelogenesis imperfecta [36], and inhibition of thoracic aortic aneurysms in mice with Marfan syndrome [30], are consistent with decreased active TGFβ levels, commensurate with a requirement for LTBP3 to target the LLC to the extracellular matrix (ECM) for latent TGFβ activation. However, it must be stated that in none of these examples has a decrease in levels of mature TGFβ in the tissue been rigorously demonstrated.

Recently Halbgebauer et al. [43] reported that knockout of LTBP3 in human Simpson-Golabi-Behmel cells and human primary adipose-derived stromal cells had no effect upon adipogenesis with respect to markers of white fat differentiation, such as PPARγ, but did affect the expression of UCP-1, a marker for brown fat. It is unclear why these results differ from the results reported here, but there are differences in cell source, differentiation medium, method of LTBP depletion, and length of time of the assay. It is important to note that he length of time of our assays were limited to 5-7 days because of the transitory nature of the KD. It will be important to clarify the explanation (s) for the differences between our results and those of Halbgebauer et al.

Although our results are contradictory to these earlier results and their interpretation indicating loss of LTBP yields decreased TGFβ, our findings are in agreement with multiple reports describing enhanced TGFβ levels after perturbation or loss of specific matrix proteins. Neptune et al. [44] reported that enhanced levels of active TGFβ accounted for aspects of Marfan syndrome caused by mutations in fibrillin1, a major partner for LTBP binding and crosslinking [45, 46]. The authors reasoned that under conditions of decreased or defective fibrillin1, LLCs were improperly sequestered in the ECM permitting inappropriate latent TGFβ activation. Recent experiments from the Ramirez laboratory demonstrating that fibrillin1 loss yields enhanced levels of active TGFβ, especially with BMSC (*vide infra*), support this interpretation [25, 47]. Heightened levels of TGFβ signaling are also observed upon perturbation of other ECM proteins, especially those that interact with fibrillin, including elastin [48, 49], ADAMTSL2 [50], MAGP1 [51, 52], LTBP4 [53], Fibulin 4 [54, 55], and proteoglycans [56,57,58]. Most recently, Abriel et al. [59] described heightened levels of TGFβ in the zebrafish outflow tract with deletions of LTBP1 and 3. Similarly, a recent report of LTBP1 with C-terminal truncation in human describes heightened TGFβ levels in cultured cells [60].

Several different mechanisms may account for the heightened TGFβ observed upon ECM protein loss. Increased active TGFβ in mice with null mutations for MAGP-1 [51, 52] or proteoglycans [56,57,58] may represent a deficiency of TGFβ binding molecules. Alternatively, enhanced TGFβ levels observed after perturbation of amount or distribution of proteins, such as fibrillin, involved in the binding of latent TGFβ complexes may reflect the misdirection and inappropriate activation of latent complexes, as originally proposed by Neptune et al. [44]. Increased TGFβ observed after the loss of elastin, LTBP4, or ADAMTSL2 [48,49,50,53] may reflect the response, i.e., increased production of the inducer of matrix protein production (TGFβ), of cells to a failed matrix [61].

However, these explanations fail to account for our observations and those of Abriel et al. [59]. The earlier results, unlike our studies, all measured TGFβ changes under conditions in which there is no reported decrease in LLC production. In our experiments active TGFβ is unlikely to derive from an LLC, as LTBP1, 2, and 4 levels decrease significantly during differentiation. However, this possibility has not yet been excluded by our in vitro experiments that are relatively short term. Alternatively, the SLC could bind to GARP or LRRC32, but these molecules are not known to be expressed by adipocytes. It is also possible that direct activators of the SLC may be present or, alternatively, the *Ltbp3* KD cells might directly release mature TGFβ. The exploration of these possible mechanisms is currently under investigation, as well as the basis for the loss of LTBP3 yielding opposite effects in different cells.

## Materials and Methods

### Mice

Generation of *Ltbp3^-/-^* mice was described previously [18]. All animal experiments were performed with approval from the Institutional Animal Care and Use Committees of the New York University Grossman School of Medicine. Mice were fed a standard chow diet (13% kcal fat, LabDiet, no. 5053) with DietGel® Boost (72-04-5022 2 oz (56 g). All experiments were performed with adult male mice, approximately 18 weeks of age.

### Body composition

Body composition (% fat mass and % lean mass) of 18-week-old, age matched *Ltbp3^-/-^* and *WT* male and female mice was assessed using a Lunar PIXImus Dual-X-ray energy absorptiometry (DEXA) instrument (Lunar Corp., Madison, WI). At the end of experiments, mice were euthanized with CO_2_, opened to visualize fat depots, and photographed. Adipose tissues, eWAT, iWAT, and liver were excised and weights determined.

### Cell lines and primary cells

C3H/10T1/2 cells were obtained from ATCC, (Manassas, VA; CCL-226), HEK293T cells from Dr. D. Bar-Sagi, M2 mouse Bone Marrow Stromal Cell (BMSC) from Dr. P. Mignatti, and 3T3-L1 cells from Dr. R. Schneider, NYU Grossman School of Medicine. BMSC were isolated as reported [62] from 8-week-old C57BL/6J male mice and cultured at 37°C with 5% CO_2_. Cells were maintained in DMEM (Corning;10-013-CV) supplemented with 10% FBS (Thermo Fisher Scientific, Gibco^TM^16140-071) and 1% Penicillin/Streptomycin (Thermo Fisher Scientific, Gibco^TM^ 15140-122). Media was changed on alternate days until the 5th day, when cells were passaged using 0.05% Trypsin-EDTA (Thermo Fisher Scientific; 25300-062), replated, and expanded for another two days before use. HEK 293T, 3T3-L1, 10T1/2, and M2 cells were subcultured every 3 days and maintained in DMEM supplemented with 10% heat-inactivated FBS and 1% Penicillin/Streptomycin.

### Adipogenic differentiation

For adipogenic differentiation experiments, BMSC cells were seeded at a density of 1 × 10^6^ cells/well in a cluster of 6 well plates. Other cells were seeded at a density of ∼3 × 10^3^ cells/cm^2^ in 6 well cluster plates. At >95% confluency, cultures were changed to mouse adipogenic differentiation media (Stem Cell Technology, Vancouver Canada; 05507), allowed to differentiate, and either harvested or fixed with 4% PFA at the times specified.

### RNA silencing

siRNA transfections were carried out using 20 pmol of siRNA (**SI Table 1**) targeting LTBP3 (siLtbp3-1-4) and lipofectamine RNAiMAX (Invitrogen, Waltham, MA; 13778-075) according to the manufacturer’s protocol. Cells were assayed on days 3 and 5 post initiation of adipogenic differentiation for gene expression at mRNA and protein levels.

### Quantification of cell number

For quantification of total cell number for the computation of the percent cells with lipid droplets or vesicles, on day 5 post adipogenesis initiation, cells were fixed with 4% PFA and washed 3 times with 1X PBS. DAPI (Thermo Fisher Scientific; 62248) staining (0.66 µg/µL) was performed for 5 min at room temperature followed by washing 3 times with 1X PBS. Cells were photographed using the Bio-Red, ZOE™ Fluorescent Cell Imager. Nuclei (blue) and cells with lipid vesicles (white) were counted manually. The percentage of cells with lipid vesicles was calculated using the formula: Percentage of cells with lipid vesicles = (Number of cells with lipid droplets/Total number of nuclei) × 100.

### TGFβ inhibition or supplementation experiments

Forty-eight hours post siRNA treatment, siCtrl or siLtbp3-2 cells were exposed to the pan-TGFβ neutralizing antibody 1D11 (Bio X Cell, Lebanon, NH;1D11.16.8, BP0057; 80 ng/mL) or an isotype-matched murine IgG (13C4; gift of F. Ramirez, Mount Sinai School of Medicine) in regular media for 4 h and then in adipogenic media for 5 days with fresh medium added after 3 days. On day 5, post adipogenic treatment, cells were fixed with 4% PFA and stained with ORO. For the inhibition of TGFβ type I receptor, LTBP3 KD cells were treated with 2 μM kinase inhibitor SB431542 (Millipore Sigma; S4317-5MG) or DMSO as a vehicle control for SB431542 in basal media for 4 h followed by incubation in differentiation media supplemented with 2 μM inhibitor for 3 to 5 days. Cells were fixed using 4% PFA followed by DAPI staining and imaging as described below. To test the effect of TGFβ supplementation, cultured cells treated with siCtrl or siLTBP3-2 cells for 48 h were incubated with dosages of 0, 0.062, or 1.25 ng/mL of TGFβ1 (R & D Systems; 7346-B2-005) in adipogenic media for 5 days and assayed for mature adipocyte formation based on cells with lipid vesicles or lipid droplets.

### Construction of lentivirus (LV) expressing *Ltbp3*

Full-length *Ltbp3* cDNA (VectorBuilder; Chicago, IL) was cloned into the pLV[Exp]-EGFP:T2A:Puro-EF1A vector through Vector Builder services (VB200618-1318juf). The final plasmid was sequenced to confirm correct insertion of *Ltbp3* ORF. The lentiviral particles were produced by Vector Builder (titer ∼1x 10^8^/mL).

### LTBP3 production in vitro

To quantify in vitro LTBP3 production, we transduced cells with either pLV[Exp]-Puro-EF1A>{mLtbp3[NM_008520.3] (VB200618-1318juf) or control vector pLV[Exp]-EGFP:T2A:Puro-EF1A>mCherry (VB160109-10005) at a multiplicity of infection of 5-10. LTBP3 synthesis and secretion was measured by immunofluorescence (day 6 post lentiviral transduction) and by immunoblotting of cell lysates (day 8 post transduction).

### RNA isolation, cDNA synthesis, and quantitative RT-PCR

Total RNA was isolated using QIAzol lysis reagent (QIAGEN, Hilden, Germany; 79306) and QIAGEN Mini RNeasy kit (QIAGEN; 74004). Genomic DNA was digested using RNAse-Free DNase (QIAGEN; 79254). Equal amounts of RNA were converted into cDNA using an iScript cDNA synthesis kit (Bio-Rad, Hercules, CA; 1708891) following the manufacturer’s instructions. Quantitative Real-Time PCR for mRNA levels were measured on a 7500 FAST RT-PCR using TaqMan probe and universal advanced master mix (Thermo Fisher Scientific, TaqMan^TM^ Fast Universal PCR Master Mix (2X), AmpErase™ UNG, 4367846). Relative mRNA expression was determined by the ΔΔCt method normalized with housekeeping genes beta-2 microglobulin (*B2m*). Fold change relative mRNA expression was determined by 2^∧-ΔΔCt^ as described [62, 63]. The list of primers used is given in **SI Table 2**.

### Protein extraction and immunoblotting

Cells were lysed in RIPA lysis buffer containing 50 mM Tris pH 7.5, 150 mM NaCl, 1% NP-40, 0.5% sodium deoxycholate, 0.1% SDS, supplemented with protease (Complete, Roche) and phosphatase (PhosSTOP, Roche) inhibitor cocktails, and 1 mM phenylmethylsulfonyl fluoride (Cell Signaling, Danvers, MA; 8553). Cell lysates were cleared by centrifugation (15,000 g for 15 min at 4 °C). Protein concentrations were determined using Pierce BCA Protein Assay Kit (Thermo Scientific, 23227). Equal amounts of protein (25 μg) were separated by SDS-PAGE, and transferred onto Nitrocellulose Membranes (Bio-Rad; 1620212) using ExpressPlus™ PAGE Gel 4-20% (GeneScript, Piscataway, NJ; M42015) wet transfer (60V for 2 h). Thereafter, membranes were blocked for 1 h in intercept (PBS) blocking buffer (LI-COR Biosciences; P/N 927-70001), followed by incubation with the indicated primary antibody (LTBP-3, pAb952) [56]; PPARγ (Cell Signaling Technology; 81B8), or GAPDH (Santa Cruz SC-32233). Secondary antibody (LI-COR Biotechnologies, Lincoln, NE; 925-68070/925-32211) incubation with anti-rabbit (LI-COR, IRDye 800CW Goat anti-Rb IgG) or anti-mouse secondary antibodies (LI-COR, IRDye 680RD Goat anti-Mouse IgG). Imaging was conducted on an Image Studio™ 5.2x Odyssey CLx, (LI-COR Lincoln, Nebraska USA). Image analysis of protein bands was determined using Image Studio™ 5.2x Odyssey CLx.

### Immunofluorescence

Cells were fixed with 2% PFA for 5 min at room temperature, washed 3 times with 1X PBS, permeabilized in 0.2% Triton X-100 for 5 min, and incubated with 5% serum from the species used to generate the secondary antibody. Cells were subsequently incubated with primary antibody overnight at 4°C, washed 3X with 1X PBS, and incubated with secondary antibody for 1 h at room temperature followed by 3 washes with 1X PBS. Primary and secondary antibodies dilutions were prepared in 1X PBST. DAPI staining (0.66 µg/µL) was performed for 5 min at room temperature followed by washing in PBST. Slides were mounted using antifade reagent (Invitrogen ProLong Gold Antifade Mountant; P10144) for 3 days at room temperature and sealed with colorless nail polish. The slides were kept at – 20°C until photographed with a Nikon microscope (Nikon ECLIPSE TS100) image software NIS-Elements D5.30.05 64 bit). Scale bar, 100 µm.

### Lipid accumulation assay

Lipid accumulation in adipocytes was detected by ORO staining (Millipore Sigma, St. Louis, MO; O0625-25G) as described [64].

### Luciferase assay

The active TGFβ in *Ltbp3* KD cells was determined using luciferase reporter cells [65]. To measure active TGFβ, 10T1/2 cells were treated with siCtrl or siLtbp3-2 for 48 h, at which time reporter cells were co-cultured at a ratio of 1.5× 10^4^ siLtbp3-2 or siCtrl cells to 2× 10^3^/ reporter cells per well in a 96 well plate for 8 h, followed by addition of the pan-TGFβ neutralizing antibody 1D11 (80 ng/mL), an isotype-matched murine IgG (13C4; 80 ng/mL), TGFβ type I receptor kinase inhibitor (SB431542; 2 μM), or DMSO for 16 h in regular or adipogenic media. The luciferase assay was performed as described [65].

### Rescue assay

Four h post-seeding, cells were transduced with either LentiCtrl or LentiLTBP3 virus particles. Day 1 post-transduction, cells were trypsinized, replated, and grown for 3 more days. On day 6 post-infection, cells were reseeded in 6 well cluster plates at a density of 3 × 10^3^ cells/cm^2^. Four h post-seeding, LentiCtrl or LentiLTBP3 cells were treated overnight with 20 pmol siCtrl or siLtbp3-2. After overnight incubation, medium was changed to fresh media for a further 24 h siRNA incubation. Cells were divided into groups – 0 h, when cells were immediately processed for RNA isolation, and 72 h, when cells were processed for measurement of mRNA levels using qRT-PCR. To measure protein levels, experimental conditions were the same as above. Cells were processed for protein extraction as described. To measure mature adipocyte numbers, cells were fixed in 4% PFA on day 5 post adipogenic induction and percentage of cells with lipid vesicles was computed as defined above.

### Statistical analysis

Data are reported as means ± SEM in bar graphs. *p < 0.05, **p < 0.01, ***p < 0.001. For *in vivo* studies, minimum 5-10 animals per group and age-matched animals were used. Cell culture experiments were reproduced 2-3 times using cells at different passage numbers. Statistical significance of data was evaluated based on the experimental conditions and comparisons as defined in the figure legend and using one of the following statistical tests; one-way analysis of variance (ANOVA), two-way mixed model analysis of variance ANOVA with Tukey’s multiple comparisons test, Kruskal-Wallis test with Dunn’s multiple comparisons tests, or Mann-Whitney U test. Data were analyzed employing GraphPad Prism 9 and JMP PRO 16.

## Author contributions

KS, NS, TE and BD performed the experiments. KS and DR wrote the manuscript. KS and DR conceptualized, designed, visualized, and analyzed the data. DR supervised the study. All authors have read the manuscript.

## Acknowledgments

We thank Megha Kothari, Princi Labana, and Avinash Singh for assisting in counting cells for the adipogenic differentiation assay.

## Availability of data

All data will be available from the DRYAD database once the manuscript is accepted.

## Supplementary material

**Supplementary Figure 1.**
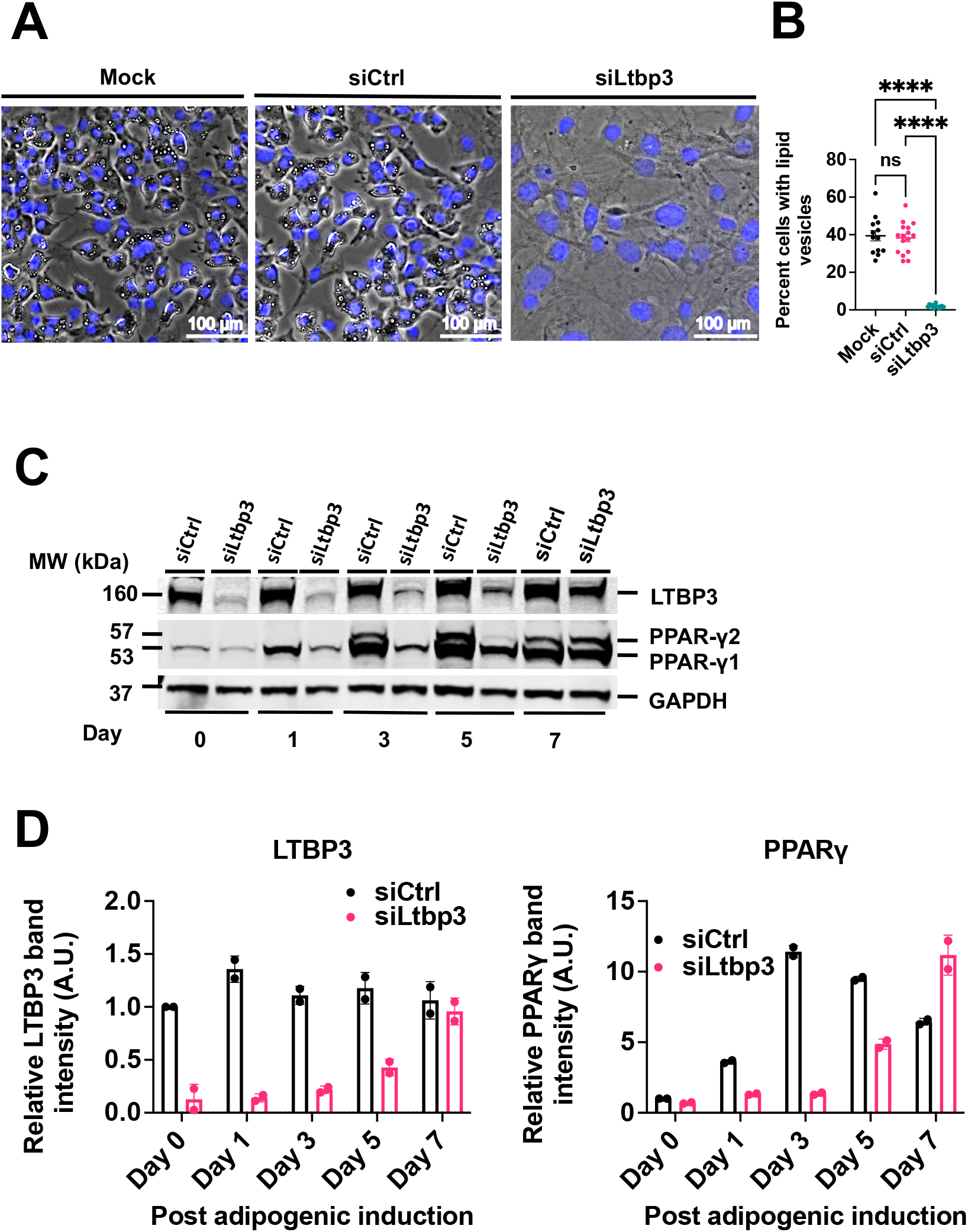
LTBP3 loss inhibits adipogenesis. **(A)** Lipid droplet accumulation after *Ltbp3* KD. Images represent siCtrl or siLtbp3-4 treated 10T1/2 cells on day 5 after adipogenic stimulation illustrating lipid droplets (white vesicles) and DAPI stained nuclei (blue). The images are representative of 3 experiments. **(B**) Quantification of lipid loaded cells from panel **A**. The number of cells with lipid droplets, as well as total number of cells, were counted manually. A total of 13-15 random fields from each treatment group were counted. **(C**) Immunoblotting for LTBP3, PPARγ, and GAPDH in 10T1/2 cells treated with siCtrl versus siLtbp3-4 for 2 days followed by adipogenic differentiation for 0, 1, 3, 5 and 7 days. The gel is representative of two independent experiments with 1 technical replica per treatment condition. **(D**) Quantification of immunoblots panel **C** for LTBP3 and PPARγ normalized with to GAPDH. Statistical significance of **Fig. B** was evaluated by nonparametric, Kruskal-Wallis test with Dunn’s multiple comparisons test. Data are represented as means ± SEM. ****p < 0.0001.

**Supplementary Figure 2.**
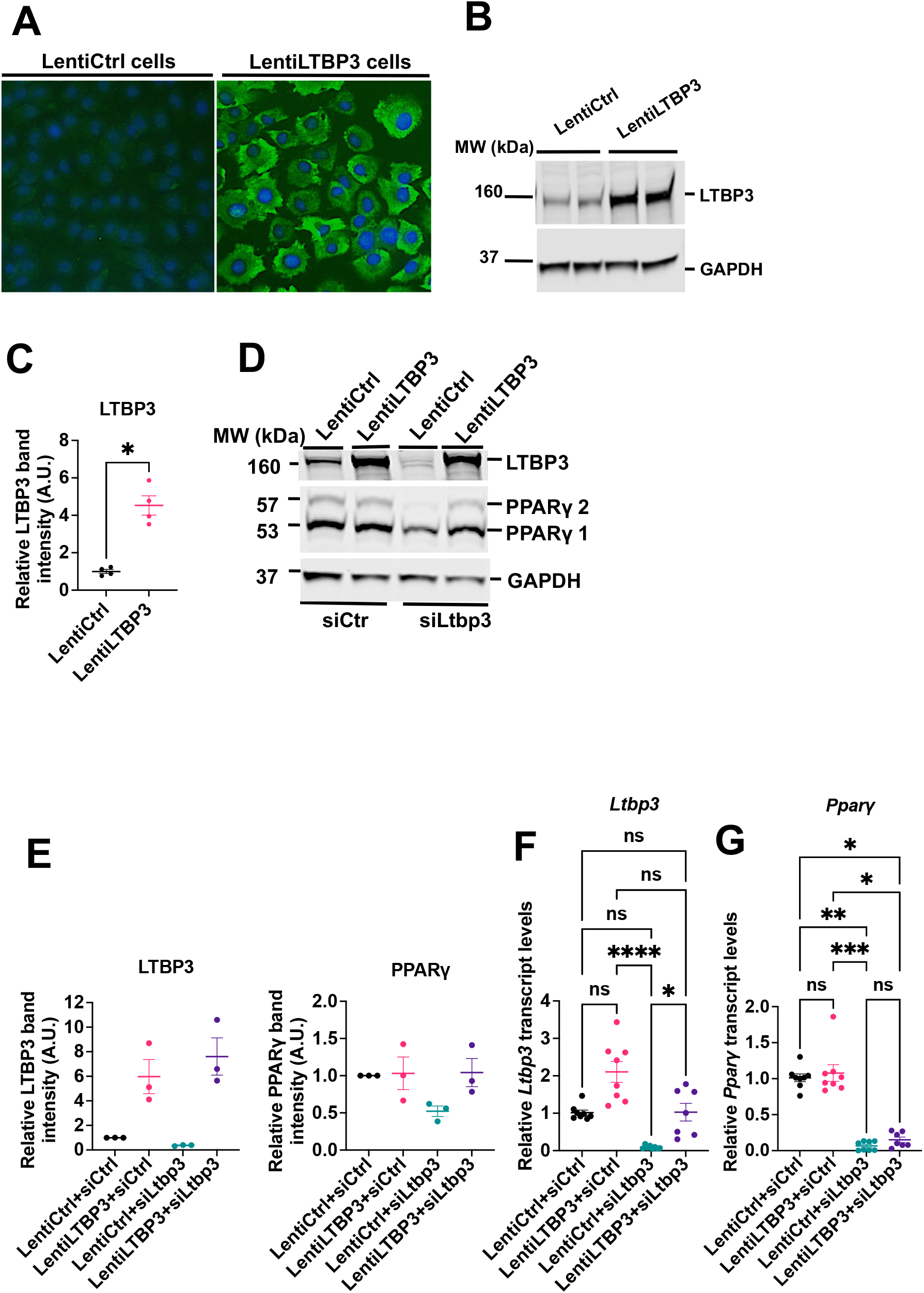
Rescue of adipogenesis by *Ltbp3* expression. (**A)** LTBP3 immunofluorescence staining. Immunostaining was performed on day 6 post exposure of 10T1/2 cells transduced with lentiviral particles expressing *Ltbp3* or control vector. Cells were fixed with 2% PFA and stained with an antibody (green) against LTBP3. Nuclei were stained with DAPI (blue). Scale bars, 100 μm. Figures are representative of one of 3 independent experiments. (**B)** Immunoblotting of LTBP3 from cell lysates after lentiviral transduction of rescue *Ltbp3* in 10T1/2 cells. The immunoblot shows the levels of LTBP3 and GAPDH in 10T1/2 transduced cells after 6 days. The immunoblot is representative of one of two independent experiments and there were two technical replicas (samples from transduction in two different wells in 6-well plate) for each group. **(C)** Quantification of immunoblots of LTBP3 normalized with GAPDH for panel **B**. There were 2 independent experiments in each group. (**D**) Immunoblots of lysates from LentiCtrl and LentiLTBP3 cells treated with siCtrl or siLtbp3-2. After 72 h of adipogenic induction, cell lysates were analyzed by immunoblotting with antibodies to LTBP3, PPARγ, and GAPDH. The immunoblot is representative of one of three independent experiments. **(E)** Quantification of immunoblots of LTBP3 and PPARγ that were normalized with GAPDH for panel **D**. The data represent the average of 3 independent experiments per treatment group and there was one technical replica for each group. **(F** and **G)** Relative mRNA levels of *Ltbp3*, and *Pparγ* in 10T1/2 cells infected with a lentivirus expressing *WT Ltbp3* and treated with siLtbp3-4 followed by 3 days of adipogenic differentiation. qRT-PCR values were normalized to *B2m* and plotted relative to siCtrl. The figures represent the average of 2 independent experiments per treatment group and there were 3-4 technical replicas for per group. Each technical replica was analyzed twice with qRT-PCR. Statistical significance of Fig. 2C was evaluated with Mann-Whitney U test, and of Fig. 2F and **G** was evaluated using nonparametric, Kruskal-Wallis test with Dunn’s multiple comparisons test. Data are represented as means ± SEM. *p < 0.05, **p < 0.01, ***p < 0.001, ****p < 0.0001.

**Supplementary Figure 3.**
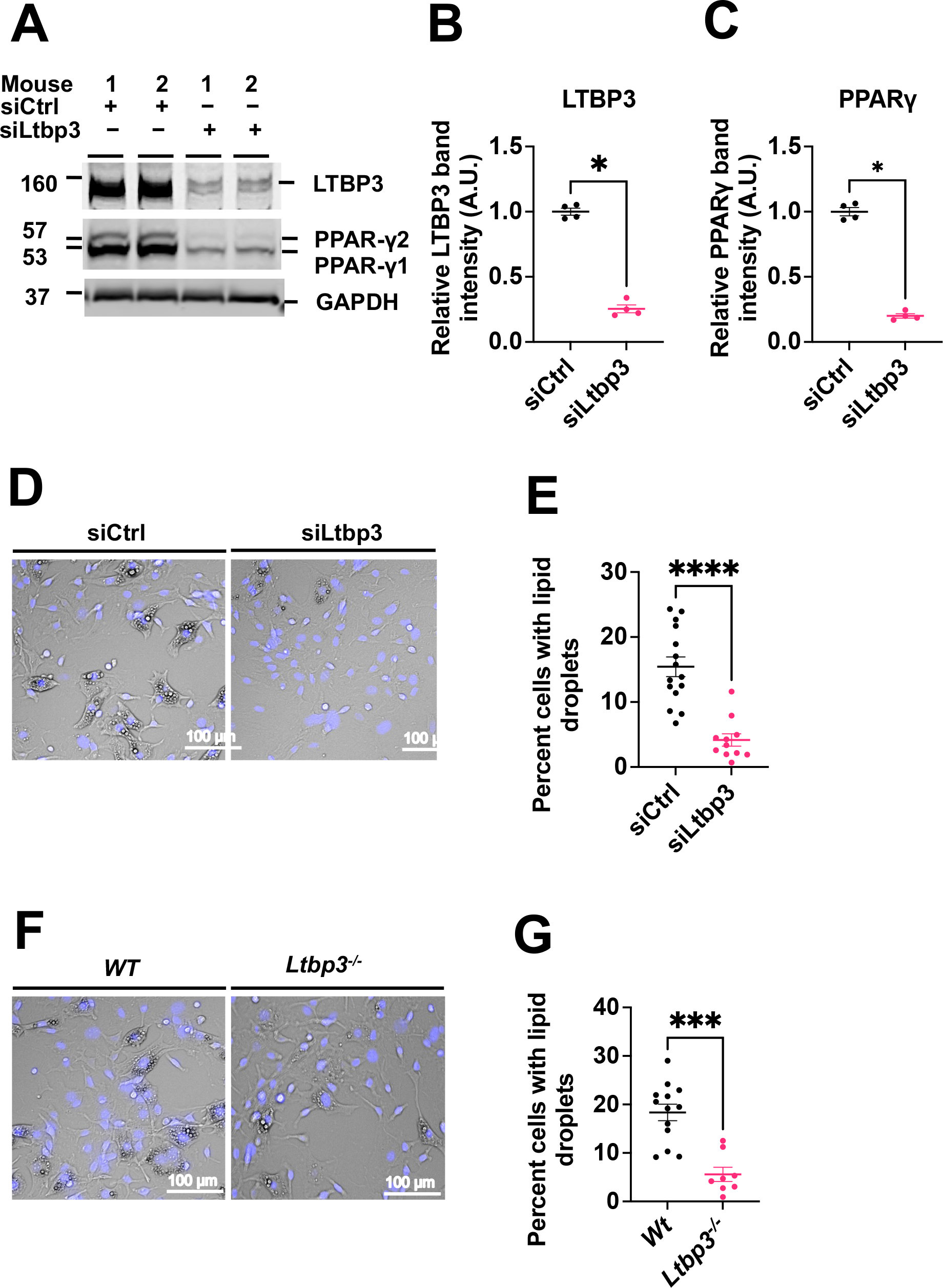
LTBP3 loss in BMSC inhibits adipogenesis. **(A)** Immunoblot illustrating the levels of LTBP3, PPARψ, and GAPDH in BMSC treated with siCtrl or siLtbp3-2 for two days followed by 3 days of adipogenic differentiation media treatment. The experiment was repeated twice with two samples in each group. (**B** and **C**) Quantification of LTBP3 and PPARγ from figure **A**. Results are representative of two independent experiments. (**D**) Representative photographs illustrating the lipid vesicles or droplets (white) and DAPI stained nuclei (blue) in BMSC treated with siCtrl or siLtbp3-2 followed by 5 days exposure to adipogenic differentiation media. Data were taken from one of two independent experiments. There were two samples for each treatment and there were 4 animals in each group. 3-4 random fields were photographed for each condition. Scale bar, 100 µm. (**E**) Quantification of lipid loaded cells from panel **D**. The number of cells with lipid droplets, as well as total number of cells, were determined manually. 12-16 random fields for each group were scored. (**F**) Representative photographs illustrating the lipid vesicles or droplets (white) and DAPI stained nuclei (blue) in BMSC from *Wt* and *Ltbp3^-/-^* mice treated with adipogenic differentiation media for 5 days. Data were taken from one of two independent experiments. There were two samples for each group. 3-4 random fields were photographed for each group. Scale bar, 100 µm. (**G**) Quantification of lipid loaded cells from panel **F.** The number of cells with lipid vesicles, as well as total number of cells, were computed manually. 8-13 random fields for each group were scored (2,000-4,000 cells for each group). Statistical significance of Fig. B, C, E, and **G** was confirmed using Mann-Whitney U test. Data are represented as means ± SEM. *p < 0.05, **p < 0.01, ***p < 0.001.

**Supplementary Figure 4.**
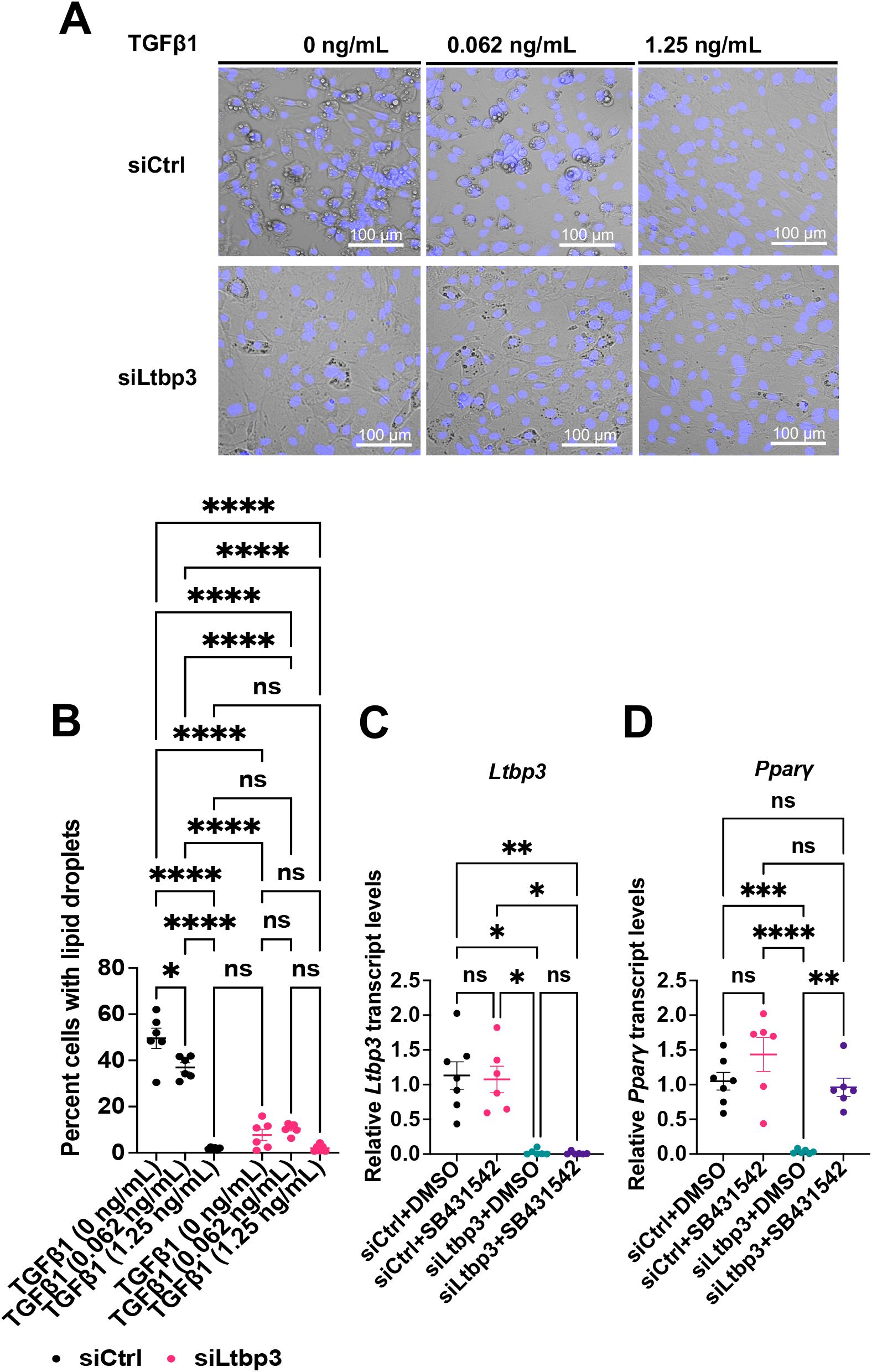
TGFβ treatment inhibits adipogenesis, whereas TGFβ receptor type I signaling inhibition rescues adipogenesis. (**A**) Representative images showing accumulation of lipid droplets (white vesicles) and DAPI stained nuclei (blue) in 10T1/2 cells treated with siCtrl or siLtbp3-2 for two days and exposed to different dosages of TGFβ1 (0, 0.062, and 1.25 ng/mL) in adipogenic media for 5 days. (**B**) Quantification of lipid loaded cells from **A**. The number of cells with lipid droplets, as well as total number of cells, were counted manually. 6 random fields from each group were scored. Data are representative of two independent experiments.There were 3 technical replicas for each treatment . Random images from 4-8 fields were scored for each condition (2,000-4000 cells for each technical replica). (**C** and **D**) Relative mRNA levels of *Ltbp3 and Pparγ* in 10T1/2 cells treated with siCtrl or siLtbp3-2 with or without TGFβ receptor I kinase inhibitor. The data are from three independent experiments with 2 technical replicas for each treatment group. Each technical replica was analyzed twice by qRT-PCR. Statistical significance was evaluated by the two-way mixed model ANOVA with Tukey’s multiple comparisons test (**B**), one-way **ANOVA** with Tukey’s multiple comparisons test (**C**) or by using nonparametric, Kruskal-Wallis test with Dunn’s multiple comparisons test (**D**). Data are represented as means ± SEM. p < 0.05, **p < 0.01, ***p < 0.001, ****p < 0.0001.

## Supplementary Tables

**Supplementary Table 1.**
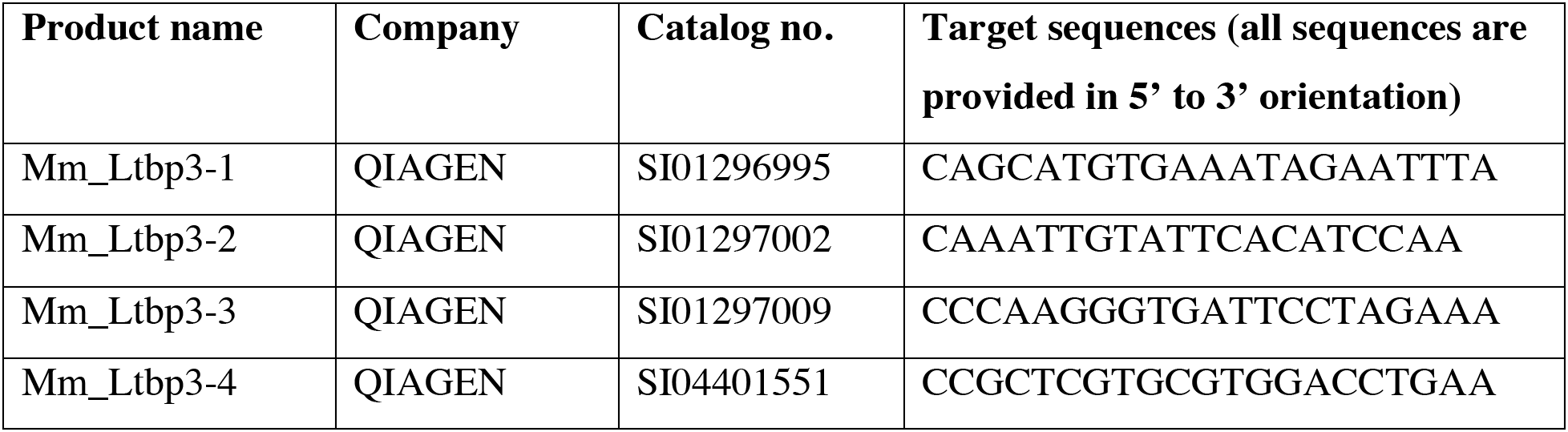
Nucleotide sequences of mouse *Ltbp3*-specific siRNAs used in the study.

**Supplementary Table 2.**
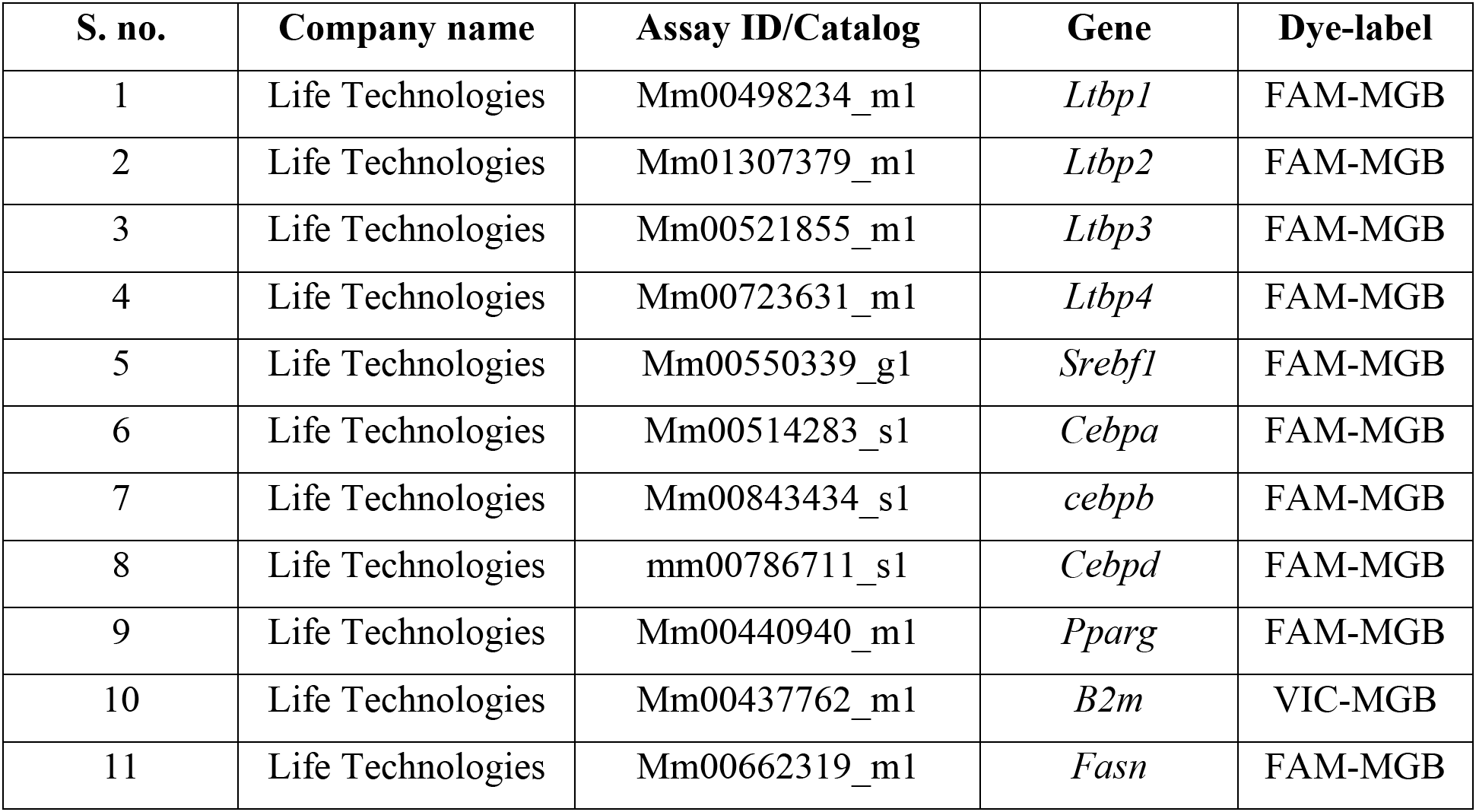
Taqman primers used in the study.

## References

1. S.J. Engle, J.B. Hoying, G.P. Boivin, I. Ormsby, P.S. Gartside, T. Doetschman, Transforming growth factor beta1 suppresses nonmetastatic colon cancer at an early stage of tumorigenesis, Cancer Res. 59 (1999) 3379–3386.

2. Z. Yang, Z. Mu, B. Dabovic, V. Jurukovski, D. Yu, J. Sung, X. Xiong, J.S. Munger, Absence of integrin-mediated TGFbeta1 activation in vivo recapitulates the phenotype of TGFbeta1-null mice, J. Cell Biol. 176 (2007) 787–793.

3. K. Yoshinaga, H. Obata, V. Jurukovski, R. Mazzieri, Y. Chen, L. Zilberberg, D. Huso, J. Melamed, P. Prijatelj, V. Todorovic, B. Dabovic, D.B. Rifkin, Perturbation of transforming growth factor (TGF)-beta1 association with latent TGF-beta binding protein yields inflammation and tumors, Proc. Natl. Acad. Sci. U. S. A. 105 (2008) 18758–18763.

4. M. Morikawa, R. Derynck, K. Miyazono, TGF-β and the TGF-β Family: Context-Dependent Roles in Cell and Tissue Physiology, Cold Spring Harb. Perspect. Biol. 8 (2016). https://doi.org/10.1101/cshperspect.a021873.

5. D.B. Constam, Regulation of TGFβ and related signals by precursor processing, Semin. Cell Dev. Biol. 32 (2014) 85–97.

6. I.B. Robertson, D.B. Rifkin, Regulation of the Bioavailability of TGF-β and TGF-β-Related Proteins, Cold Spring Harb. Perspect. Biol. 8 (2016). https://doi.org/10.1101/cshperspect.a021907.

7. S. Liénart, R. Merceron, C. Vanderaa, F. Lambert, D. Colau, J. Stockis, B. van der Woning, H. De Haard, M. Saunders, P.G. Coulie, S.N. Savvides, S. Lucas, Structural basis of latent TGF-β1 presentation and activation by GARP on human regulatory T cells, Science. 362 (2018) 952–956.

8. Y. Qin, B.S. Garrison, W. Ma, R. Wang, A. Jiang, J. Li, M. Mistry, R.T. Bronson, D. Santoro, C. Franco, D.A. Robinton, B. Stevens, D.J. Rossi, C. Lu, T.A. Springer, A Milieu Molecule for TGF-β Required for Microglia Function in the Nervous System, Cell. 174 (2018) 156–171.e16. https://doi.org/10.1016/j.cell.2018.05.027.

9. T. Harel, E. Levy-Lahad, M. Daana, H. Mechoulam, S. Horowitz-Cederboim, M. Gur, V. Meiner, O. Elpeleg, Homozygous stop-gain variant in LRRC32, encoding a TGFβ receptor, associated with cleft palate, proliferative retinopathy, and developmental delay, Eur. J. Hum. Genet. 27 (2019) 1315–1319.

10. D. Rifkin, N. Sachan, K. Singh, E. Sauber, G. Tellides, F. Ramirez, The role of LTBPs in TGF beta signaling, Dev. Dyn. 251 (2022) 95–104.

11. J. Saharinen, J. Taipale, J. Keski-Oja, Association of the small latent transforming growth factor-beta with an eight cysteine repeat of its binding protein LTBP-1, EMBO J. 15 (1996) 245–253.

12. P.-E. Gleizes, R.C. Beavis, R. Mazzieri, B. Shen, D.B. Rifkin, Identification and Characterization of an Eight-cysteine Repeat of the Latent Transforming Growth Factor-β Binding Protein-1 that Mediates Bonding to the Latent Transforming Growth Factor-β1, Journal of Biological Chemistry. 271 (1996) 29891–29896. https://doi.org/10.1074/jbc.271.47.29891.

13. J. Saharinen, J. Keski-Oja, Specific sequence motif of 8-Cys repeats of TGF-beta binding proteins, LTBPs, creates a hydrophobic interaction surface for binding of small latent TGF-beta, Mol. Biol. Cell. 11 (2000) 2691–2704.

14. D.B. Rifkin, Latent transforming growth factor-beta (TGF-beta) binding proteins: orchestrators of TGF-beta availability, J. Biol. Chem. 280 (2005) 7409–7412.

15. D.E. Clouthier, S.A. Comerford, R.E. Hammer, Hepatic fibrosis, glomerulosclerosis, and a lipodystrophy-like syndrome in PEPCK-TGF-beta1 transgenic mice, Journal of Clinical Investigation. 100 (1997) 2697–2713. https://doi.org/10.1172/jci119815.

16. L. Choy, J. Skillington, R. Derynck, Roles of autocrine TGF-beta receptor and Smad signaling in adipocyte differentiation, J. Cell Biol. 149 (2000) 667–682.

17. L. Choy, R. Derynck, Transforming growth factor-beta inhibits adipocyte differentiation by Smad3 interacting with CCAAT/enhancer-binding protein (C/EBP) and repressing C/EBP transactivation function, J. Biol. Chem. 278 (2003) 9609–9619.

18. B. Dabovic, Y. Chen, C. Colarossi, H. Obata, L. Zambuto, M.A. Perle, D.B. Rifkin, Bone abnormalities in latent TGF-β binding protein (Ltbp)-3–null mice indicate a role for Ltbp-3 in modulating TGF-β bioavailability, Journal of Cell Biology. 156 (2002) 227–232. https://doi.org/10.1083/jcb.200111080.

19. K. Koli, M. Hyytiäinen, M.J. Ryynänen, J. Keski-Oja, Sequential deposition of latent TGF-beta binding proteins (LTBPs) during formation of the extracellular matrix in human lung fibroblasts, Exp. Cell Res. 310 (2005) 370–382.

20. R.A. Ignotz, J. Massagué, Type beta transforming growth factor controls the adipogenic differentiation of 3T3 fibroblasts, Proc. Natl. Acad. Sci. U. S. A. 82 (1985) 8530–8534.

21. R.L. Sparks, R.E. Scott, Transforming growth factor type β is a specific inhibitor of 3T3 T mesenchymal stem cell differentiation, Experimental Cell Research. 165 (1986) 345–352. https://doi.org/10.1016/0014-4827(86)90588-4.

22. F.M. Torti, S.V. Torti, J.W. Larrick, G.M. Ringold, Modulation of adipocyte differentiation by tumor necrosis factor and transforming growth factor beta, J. Cell Biol. 108 (1989) 1105–1113.

23. E.J. van Zoelen, I. Duarte, J.M. Hendriks, S.P. van der Woning, TGFβ-induced switch from adipogenic to osteogenic differentiation of human mesenchymal stem cells: identification of drug targets for prevention of fat cell differentiation, Stem Cell Res. Ther. 7 (2016) 123.

24. N. Zamani, C.W. Brown, Emerging roles for the transforming growth factor-{beta} superfamily in regulating adiposity and energy expenditure, Endocr. Rev. 32 (2011) 387–403.

25. S. Smaldone, N.P. Clayton, M. del Solar, G. Pascual, S.H. Cheng, B.M. Wentworth, M.B. Schaffler, F. Ramirez, Fibrillin-1 Regulates Skeletal Stem Cell Differentiation by Modulating TGFβ Activity Within the Marrow Niche, J. Bone Miner. Res. 31 (2016) 86–97.

26. R. Serra, M. Johnson, E.H. Filvaroff, J. LaBorde, D.M. Sheehan, R. Derynck, H.L. Moses, Expression of a truncated, kinase-defective TGF-beta type II receptor in mouse skeletal tissue promotes terminal chondrocyte differentiation and osteoarthritis, J. Cell Biol. 139 (1997) 541–552.

27. M.B. Mueller, M. Fischer, J. Zellner, A. Berner, T. Dienstknecht, L. Prantl, R. Kujat, M. Nerlich, R.S. Tuan, P. Angele, Hypertrophy in mesenchymal stem cell chondrogenesis: effect of TGF-beta isoforms and chondrogenic conditioning, Cells Tissues Organs. 192 (2010) 158–166.

28. I. Grafe, S. Alexander, J.R. Peterson, T.N. Snider, B. Levi, B. Lee, Y. Mishina, TGF-β Family Signaling in Mesenchymal Differentiation, Cold Spring Harb. Perspect. Biol. 10 (2018). https://doi.org/10.1101/cshperspect.a022202.

29. M.L. Muthu, K. Tiedemann, J. Fradette, S. Komarova, D.P. Reinhardt, Fibrillin-1 regulates white adipose tissue development, homeostasis, and function, Matrix Biol. 110 (2022) 106–128.

30. L. Zilberberg, C.K. Phoon, I. Robertson, B. Dabovic, F. Ramirez, D.B. Rifkin, Genetic analysis of the contribution of LTBP-3 to thoracic aneurysm in Marfan syndrome, Proc Natl Acad Sci U S A. 112 (2015) 14012–7. doi: 10.1073/pnas.1507652112 Epub 2015 Oct 22. PMID: 26494287; PMCID: PMC4653215.

31. B. Dabovic, Y. Chen, J. Choi, E.C. Davis, L.Y. Sakai, V. Todorovic, M. Vassallo, L. Zilberberg, A. Singh, D.B. Rifkin, Control of lung development by latent TGF-βbinding proteins, J. Cell. Physiol. 226 (2011) 1499–1509.

32. K. Koli, M.J. Ryynänen, J. Keski-Oja, Latent TGF-beta binding proteins (LTBPs)-1 and -3 coordinate proliferation and osteogenic differentiation of human mesenchymal stem cells, Bone. 43 (2008) 679–688.

33. A. Gualandris, J.P. Annes, M. Arese, I. Noguera, V. Jurukovski, D.B. Rifkin, The latent transforming growth factor-beta-binding protein-1 promotes in vitro differentiation of embryonic stem cells into endothelium, Mol. Biol. Cell. 11 (2000) 4295–4308.

34. Y. Nakajima, K. Miyazono, M. Kato, M. Takase, T. Yamagishi, H. Nakamura, Extracellular fibrillar structure of latent TGF beta binding protein-1: role in TGF beta-dependent endothelial-mesenchymal transformation during endocardial cushion tissue formation in mouse embryonic heart, J. Cell Biol. 136 (1997) 193– 204.

35. R. Flaumenhaft, M. Abe, Y. Sato, K. Miyazono, J. Harpel, C.H. Heldin, D.B. Rifkin, Role of the latent TGF-beta binding protein in the activation of latent TGF-beta by co-cultures of endothelial and smooth muscle cells, J. Cell Biol. 120 (1993) 995–1002.

36. M. Huckert, C. Stoetzel, S. Morkmued, V. Laugel-Haushalter, V. Geoffroy, J. Muller, F. Clauss, M.K. Prasad, F. Obry, J.L. Raymond, M. Switala, Y. Alembik, S. Soskin, E. Mathieu, J. Hemmerlé, J.-L. Weickert, B.B. Dabovic, D.B. Rifkin, A. Dheedene, E. Boudin, O. Caluseriu, M.-C. Cholette, R. Mcleod, R. Antequera, M.-P. Gellé, J.-L. Coeuriot, L.-F. Jacquelin, I. Bailleul-Forestier, M.-C. Manière, W. Van Hul, D. Bertola, P. Dollé, A. Verloes, G. Mortier, H. Dollfus, A. Bloch-Zupan, Mutations in the latent TGF-beta binding protein 3 (LTBP3) gene cause brachyolmia with amelogenesis imperfecta, Hum. Mol. Genet. 24 (2015) 3038– 3049.

37. M. Horiguchi, V. Todorovic, K. Hadjiolova, R. Weiskirchen, D.B. Rifkin, Abrogation of both short and long forms of latent transforming growth factor-β binding protein-1 causes defective cardiovascular development and is perinatally lethal, Matrix Biology. 43 (2015) 61–70. https://doi.org/10.1016/j.matbio.2015.03.006.

38. J.S. Munger, X. Huang, H. Kawakatsu, M.J. Griffiths, S.L. Dalton, J. Wu, J.F. Pittet, N. Kaminski, C. Garat, M.A. Matthay, D.B. Rifkin, D. Sheppard, The integrin alpha v beta 6 binds and activates latent TGF beta 1: a mechanism for regulating pulmonary inflammation and fibrosis, Cell. 96 (1999) 319–328.

39. J.P. Annes, Y. Chen, J.S. Munger, D.B. Rifkin, Integrin αVβ6-mediated activation of latent TGF-β requires the latent TGF-β binding protein-1, Journal of Cell Biology. 165 (2004) 723–734. https://doi.org/10.1083/jcb.200312172.

40. R. Wang, J. Zhu, X. Dong, M. Shi, C. Lu, T.A. Springer, GARP regulates the bioavailability and activation of TGFβ, Mol. Biol. Cell. 23 (2012) 1129–1139.

41. M. Shi, J. Zhu, R. Wang, X. Chen, L. Mi, T. Walz, T.A. Springer, Latent TGF-β structure and activation, Nature. 474 (2011) 343–349.

42. X. Dong, B. Zhao, R.E. Iacob, J. Zhu, A.C. Koksal, C. Lu, J.R. Engen, T.A. Springer, Force interacts with macromolecular structure in activation of TGF-β, Nature. 542 (2017) 55–59.

43. D. Halbgebauer, J. Roos, J.B. Funcke, H. Neubauer, B.S. Hamilton, E. Simon, E.Z. Amri, K.M. Debatin, M. Wabitsch, P. Fischer-Posovszky, D. Tews, Latent TGFβ-binding proteins regulate UCP1 expression and function via TGFβ2, Mol Metab. 53 (2021) 101336.

44. E.R. Neptune, P.A. Frischmeyer, D.E. Arking, L. Myers, T.E. Bunton, B. Gayraud, F. Ramirez, L.Y. Sakai, H.C. Dietz, Dysregulation of TGF-beta activation contributes to pathogenesis in Marfan syndrome, Nat. Genet. 33 (2003) 407–411.

45. Z. Isogai, R.N. Ono, S. Ushiro, D.R. Keene, Y. Chen, R. Mazzieri, N.L. Charbonneau, D.P. Reinhardt, D.B. Rifkin, L.Y. Sakai, Latent transforming growth factor beta-binding protein 1 interacts with fibrillin and is a microfibril-associated protein, J. Biol. Chem. 278 (2003) 2750–2757.

46. T. Massam-Wu, M. Chiu, R. Choudhury, S.S. Chaudhry, A.K. Baldwin, A. McGovern, C. Baldock, C.A. Shuttleworth, C.M. Kielty, Assembly of fibrillin microfibrils governs extracellular deposition of latent TGF beta, J. Cell Sci. 123 (2010) 3006–3018.

47. H. Nistala, S. Lee-Arteaga, S. Smaldone, G. Siciliano, F. Ramirez, Extracellular microfibrils control osteoblast-supported osteoclastogenesis by restricting TGF{beta} stimulation of RANKL production, J. Biol. Chem. 285 (2010) 34126– 34133.

48. Q. Hu, A. Shifren, C. Sens, J. Choi, Z. Szabo, B.C. Starcher, R.H. Knutsen, J.M. Shipley, E.C. Davis, R.P. Mecham, Z. Urban, Mechanisms of emphysema in autosomal dominant cutis laxa, Matrix Biol. 29 (2010) 621–628.

49. B. Callewaert, M. Renard, V. Hucthagowder, B. Albrecht, I. Hausser, E. Blair, C. Dias, A. Albino, H. Wachi, F. Sato, R.P. Mecham, B. Loeys, P.J. Coucke, A. De Paepe, Z. Urban, New insights into the pathogenesis of autosomal-dominant cutis laxa with report of five ELN mutations, Hum. Mutat. 32 (2011) 445–455.

50. C. Le Goff, F. Morice-Picard, N. Dagoneau, L.W. Wang, C. Perrot, Y.J. Crow, F. Bauer, E. Flori, C. Prost-Squarcioni, D. Krakow, G. Ge, D.S. Greenspan, D. Bonnet, M. Le Merrer, A. Munnich, S.S. Apte, V. Cormier-Daire, ADAMTSL2 mutations in geleophysic dysplasia demonstrate a role for ADAMTS-like proteins in TGF-beta bioavailability regulation, Nat. Genet. 40 (2008) 1119–1123.

51. C.S. Craft, T.A. Pietka, T. Schappe, T. Coleman, M.D. Combs, S. Klein, N.A. Abumrad, R.P. Mecham, The extracellular matrix protein MAGP1 supports thermogenesis and protects against obesity and diabetes through regulation of TGF-β, Diabetes. 63 (2014) 1920–1932.

52. C.S. Craft, T.J. Broekelmann, W. Zou, J.C. Chappel, S.L. Teitelbaum, R.P. Mecham, Oophorectomy-induced bone loss is attenuated in MAGP1-deficient mice, J. Cell. Biochem. 113 (2012) 93–99.

53. Z. Urban, V. Hucthagowder, N. Schürmann, V. Todorovic, L. Zilberberg, J. Choi, C. Sens, C.W. Brown, R.D. Clark, K.E. Holland, M. Marble, L.Y. Sakai, B. Dabovic, D.B. Rifkin, E.C. Davis, Mutations in LTBP4 Cause a Syndrome of Impaired Pulmonary, Gastrointestinal, Genitourinary, Musculoskeletal, and Dermal Development, The American Journal of Human Genetics. 85 (2009) 593– 605. https://doi.org/10.1016/j.ajhg.2009.09.013.

54. K. Hanada, M. Vermeij, G.A. Garinis, M.C. de Waard, M.G.S. Kunen, L. Myers, A. Maas, D.J. Duncker, C. Meijers, H.C. Dietz, R. Kanaar, J. Essers, Perturbations of vascular homeostasis and aortic valve abnormalities in fibulin-4 deficient mice, Circ. Res. 100 (2007) 738–746.

55. M. Renard, T. Holm, R. Veith, B.L. Callewaert, L.C. Adès, O. Baspinar, A. Pickart, M. Dasouki, J. Hoyer, A. Rauch, P. Trapane, M.G. Earing, P.J. Coucke, L.Y. Sakai, H.C. Dietz, A.M. De Paepe, B.L. Loeys, Altered TGFβ signaling and cardiovascular manifestations in patients with autosomal recessive cutis laxa type I caused by fibulin-4 deficiency, European Journal of Human Genetics. 18 (2010) 895–901. https://doi.org/10.1038/ejhg.2010.45.

56. A. Hildebrand, M. Romarís, L.M. Rasmussen, D. Heinegård, D.R. Twardzik, W.A. Border, E. Ruoslahti, Interaction of the small interstitial proteoglycans biglycan, decorin and fibromodulin with transforming growth factor beta, Biochem. J. 302 (Pt 2) (1994) 527–534.

57. C. Cabello-Verrugio, E. Brandan, A novel modulatory mechanism of transforming growth factor-beta signaling through decorin and LRP-1, J. Biol. Chem. 282 (2007) 18842–18850.

58. H. Kizawa, I. Kou, A. Iida, A. Sudo, Y. Miyamoto, A. Fukuda, A. Mabuchi, A. Kotani, A. Kawakami, S. Yamamoto, A. Uchida, K. Nakamura, K. Notoya, Y. Nakamura, S. Ikegawa, An aspartic acid repeat polymorphism in asporin inhibits chondrogenesis and increases susceptibility to osteoarthritis, Nat. Genet. 37 (2005) 138–144.

59. M. Abrial, S. Basu, M. Huang, V. Butty, A. Schwertner, S. Jeffrey, D. Jordan, C.E. Burns, C.G. Burns, Latent TGFβ-binding proteins 1 and 3 protect the larval zebrafish outflow tract from aneurysmal dilatation, Dis. Model. Mech. 15 (2022). https://doi.org/10.1242/dmm.046979.

60. L. Pottie, C.S. Adamo, A. Beyens, S. Lütke, P. Tapaneeyaphan, A. De Clercq, P.L. Salmon, R. De Rycke, A. Gezdirici, E.Y. Gulec, N. Khan, J.E. Urquhart, W.G. Newman, K. Metcalfe, S. Efthymiou, R. Maroofian, N. Anwar, S. Maqbool, F. Rahman, I. Altweijri, M. Alsaleh, S.M. Abdullah, M. Al-Owain, M. Hashem, H. Houlden, F.S. Alkuraya, P. Sips, G. Sengle, B. Callewaert, Bi-allelic premature truncating variants in LTBP1 cause cutis laxa syndrome, Am. J. Hum. Genet. 108 (2021) 1095–1114.

61. M. Horiguchi, M. Ota, D.B. Rifkin, Matrix control of transforming growth factor-β function, J. Biochem. 152 (2012) 321–329.

62. E.M.S. Litwinoff, M.Y. Gold, K. Singh, J. Hu, H. Li, K. Cadwell, A.M. Schmidt, Myeloid ATG16L1 does not affect adipose tissue inflammation or body mass in mice fed high fat diet, Obesity Research & Clinical Practice. 12 (2018) 174–186. https://doi.org/10.1016/j.orcp.2017.10.006.

63. K. Singh, N.G. Prasad, Cold stress upregulates the expression of heat shock proteins and Frost genes, but evolution of cold stress resistance is apparently not mediated through either heat shock proteins or Frost genes in the cold stress selected population. bioRxiv. (2022). https://doi.org/10.1101/2022.03.07.483305

64. J.L. Ramírez-Zacarías, F. Castro-Muñozledo, W. Kuri-Harcuch, Quantitation of adipose conversion and triglycerides by staining intracytoplasmic lipids with Oil red O, Histochemistry. 97 (1992) 493–497.

65. M. Abe, J.G. Harpel, C.N. Metz, I. Nunes, D.J. Loskutoff, D.B. Rifkin. An assay for transforming growth factor-beta using cells transfected with a plasminogen activator inhibitor-1 promoter-luciferase construct. Anal Biochem. 216 (1994) 276–84. https://doi.org/10.1006/abio.1994.1042

